# Control of Leaf Vein Patterning by Regulated Plasmodesma Aperture

**DOI:** 10.1101/2021.12.06.471439

**Authors:** Nguyen Manh Linh, Enrico Scarpella

## Abstract

To form tissue networks, animal cells migrate and interact through proteins protruding from their plasma membranes. Plant cells can do neither, yet plants form vein networks. How plants do so is unclear, but veins are thought to form by the coordinated action of the polar transport and signal transduction of the plant hormone auxin. However, plants inhibited in both pathways still form veins. Patterning of vascular cells into veins is instead prevented in mutants lacking the function of the *GNOM* (*GN*) regulator of auxin transport and signaling, suggesting the existence of at least one more *GN*-dependent vein-patterning pathway. Here we show that pathway depends on the movement of an auxin signal through plasmodesmata (PDs) intercellular channels. PD permeability is high where veins are forming, lowers between veins and nonvascular tissues, but remains high between vein cells. Impaired ability to regulate PD aperture leads to defects in auxin transport and signaling, ultimately leading to vein patterning defects that are enhanced by inhibition of auxin transport or signaling. *GN* controls PD aperture regulation, and simultaneous inhibition of auxin signaling, auxin transport, and regulated PD aperture phenocopies null *gn* mutants. Therefore, veins are patterned by the coordinated action of three *GN*-dependent pathways: auxin signaling, polar auxin transport, and movement of an auxin signal through PDs. We have identified all the key vein-patterning pathways in plants and an unprecedented mechanism of tissue network formation in multicellular organisms.

## INTRODUCTION

The problem of long-distance transport of water, nutrients, and signaling molecules is solved in most multicellular organisms by tissue networks such as the vascular system of vertebrate embryos and the vein network of plant leaves. How tissue networks form is therefore a key question in developmental biology. In vertebrates, for example, formation of the embryonic vascular system involves direct cell–cell interaction and cell migration (reviewed, for example, in (Betz et al., 2016; Hogan and Schulte-Merker, 2017)). Both those processes are precluded in plants by a cell wall that keeps plant cells apart and in place. Therefore, leaf veins and their networks form by a different mechanism.

How leaf veins form is poorly understood, but the cell-to-cell, polar transport of the plant signaling molecule auxin has long been thought to be both necessary and sufficient for vein formation (recently reviewed in (Cieslak et al., 2021; Lavania et al., 2021)). Inconsistent with that notion, however, veins still form in mutants lacking the function of PIN-FORMED (PIN) auxin exporters, whose polar localization at the plasma membrane determines the polarity of auxin transport (Petrasek et al., 2006; Wisniewska et al., 2006; Zourelidou et al., 2014; Verna et al., 2019). By contrast, patterning of vascular cells into veins is prevented in mutants lacking the function of the guanine exchange factor for ADP ribosylation factors GNOM (GN): the vascular system of null *gn* mutants is no more than a shapeless cluster of randomly oriented vascular cells (Mayer et al., 1993; Steinmann et al., 1999; Koizumi et al., 2000; Geldner et al., 2004; Verna et al., 2019).

For more than 20 years, the vesicle trafficking regulator GN has been thought to perform its essential vein-patterning function solely through its ability to control the polarity of PIN protein localization (recently reviewed in (Lavania et al., 2021)). However, two pieces of evidence argue against that notion. First, the vein patterning defects of *gn* mutants are quantitatively stronger than and qualitatively different from those of *pin* mutants (Verna et al., 2019). Second, *pin* mutations are inconsequential to the *gn* vascular phenotype (Verna et al., 2019). These observations suggest that other pathways in addition to polar auxin transport are involved in vein patterning and that *GN* controls such additional pathways too. One such pathway is that which transduces auxin signals (Lavy and Estelle, 2016; Verna et al., 2019; Ramos Báez and Nemhauser, 2021). However, mutants impaired in both auxin signaling and polar auxin transport only phenocopy intermediate *gn* mutants (Verna et al., 2019), suggesting that additional *GN*-dependent pathways are involved in vein patterning.

Because experimental evidence suggests that auxin can move through plasmodesmata (PDs) intercellular channels (recently reviewed in (Paterlini, 2020; Band, 2021)), here we ask whether movement of auxin or an auxin-dependent signal through PDs is one of the missing *GN*-dependent vein-patterning pathways. We find that veins are patterned by the coordinated action of three *GN*-dependent pathways: auxin signaling, polar auxin transport, and movement of auxin or an auxin-dependent signal through PDs. In the most severe cases, the vascular system of leaves in which those three pathways have been inhibited is no more than a shapeless cluster of vascular cells, suggesting that we have identified all the main pathways in vein patterning.

## RESULTS

Available evidence suggests that auxin or an auxin-dependent signal (hereafter collectively referred to as an “auxin signal”) moves during leaf development, that such movement is not mediated by known auxin transporters, and that such auxin-transporter-unmediated movement controls vein patterning (Verna et al., 2019). Here we tested the hypothesis that the movement of an auxin signal that controls vein patterning and that is not mediated by auxin transporters is mediated by PDs.

### Control of Vein Patterning by Regulated PD Aperture

Should the movement of an auxin signal that controls vein patterning be mediated by PDs, defects in PD aperture regulation would lead to vein pattern defects. Because severe defects in the ability to regulate PD aperture lead to embryo lethality (e.g., (Patton et al., 1991; Kim et al., 2002; Yamagishi et al., 2005; Kobayashi et al., 2007; Stonebloom et al., 2009; Xu et al., 2012)), to test the prediction that defects in PD aperture regulation will lead to vein pattern defects, we analyzed vein patterns in mature first leaves of the *callose synthase 3 - dominant* (*cals3-d*) and *glucan-synthase-like 8* / *chorus* / *enlarged tetrad 2* / *massue* / *ectopic expression of seed storage proteins 8* (*gsl8* hereafter) mutants of Arabidopsis, which have respectively near-constitutively narrow and near-constitutively wide PD aperture and can survive embryogenesis (Chen et al., 2009; Thiele et al., 2009; Guseman et al., 2010; Vaten et al., 2011; De Storme et al., 2013; Han et al., 2014; Saatian et al., 2018).

WT Arabidopsis forms broad leaves whose vein networks are defined by at least four reproducible features: (1) a narrow I-shaped midvein that runs the length of the leaf; (2) lateral veins that branch from the midvein and join distal veins to form closed loops; (3) minor veins that branch from midvein and loops and either end freely or join other veins; and (4) minor veins and loops that curve near the leaf margin and give the vein network a scalloped outline (Telfer and Poethig, 1994; Nelson and Dengler, 1997; Kinsman and Pyke, 1998; Candela et al., 1999; Mattsson et al., 1999; Sieburth, 1999; Steynen and Schultz, 2003; Sawchuk et al., 2013; Verna et al., 2015; Verna et al., 2019) (Fig. 1A,B,F). Within individual veins, vascular elements are connected end to end and are aligned along the length of the vein, and free vein ends are as narrow as the rest of the vein (Fig. 1G).

**Figure 1.**
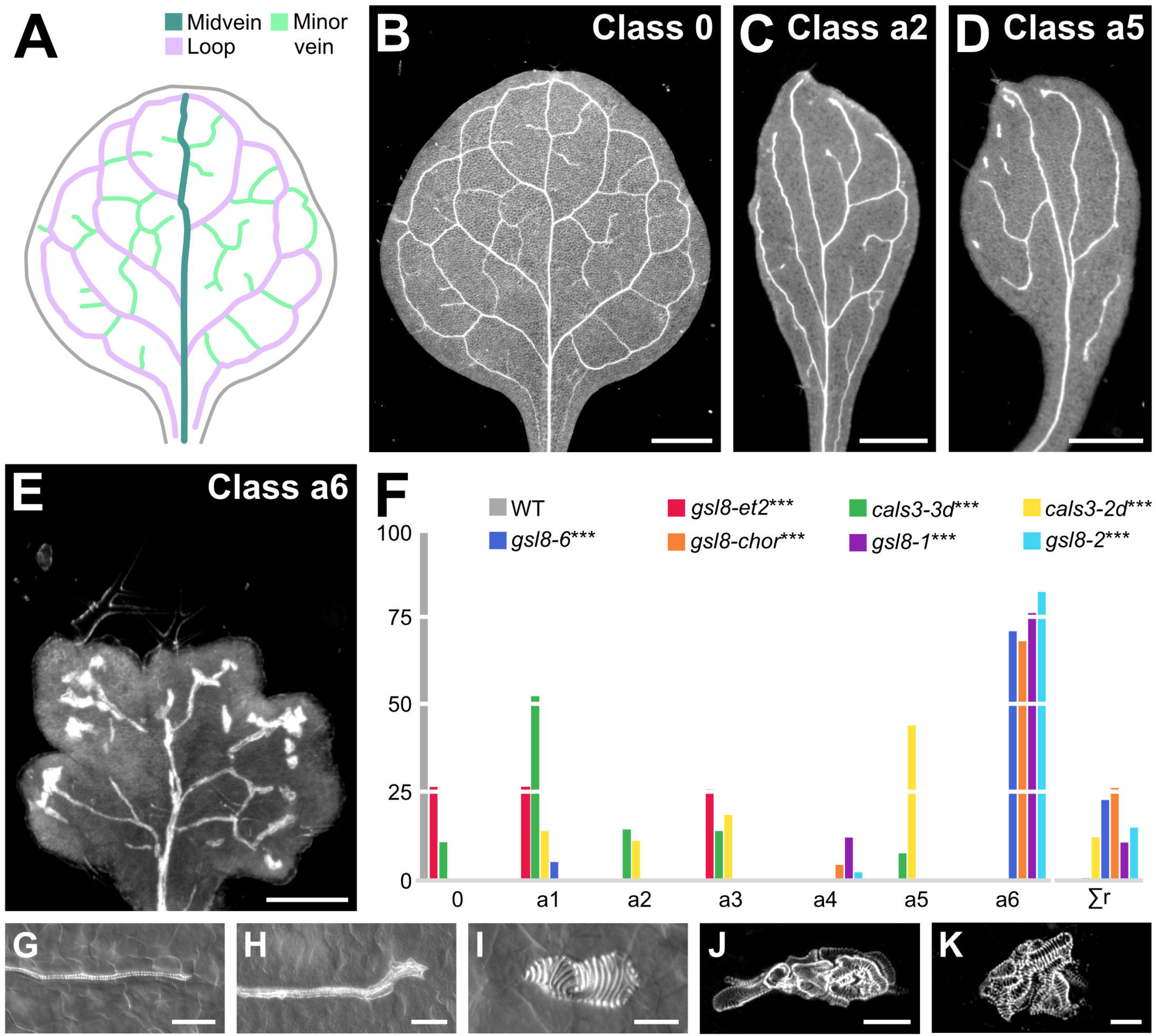
Control of Vein Patterning by PD Aperture. (A,B) Vein pattern of mature first leaf of WT Arabidopsis. In A: teal, midvein; lavender, loops; mint, minor veins. (B–E) Dark-field illumination of mature first leaves illustrating phenotype classes (top right). Class 0: narrow I-shaped midvein and scalloped vein-network outline (B). Class a2: narrow leaf and open vein-network outline (C). Class a5: narrow leaf, open vein-network outline, vein fragments, and/or vascular clusters (D). Class a6: lobed leaf, open vein-network outline, vein fragments, and/or vascular clusters (E). (F) Percentages of leaves in phenotype classes. Class a1: open vein-network outline; class a3: vein fragments and/or vascular clusters; class a4: lobed leaf, vein fragments, and/or vascular clusters (not shown). Rare vein pattern defects were grouped in class Σr. Difference between *gsl8-et2* and WT, between *cals3-3d* and WT, between *cals3-2d* and WT, between *gsl8-6* and WT, between *gsl8-chor* and WT, between *gsl8-1* and WT, and between *gsl8-2* and WT was significant at *P*<0.001 (***) by Kruskal-Wallis and Mann-Whitney test with Bonferroni correction. Sample population sizes: WT, 30; *gsl8-et2*, 108; *cals3-3d*, 216; *cals3-2d*, 173; *gsl8-6*, 39; *gsl8-chor*, 45; *gsl8-1*, 65; *gsl8-2*, 47. (G–K) Details of veins and vein ends in WT (G) and *cals3-2d* (H) or of vascular clusters in *cals3-2d* (I) and *gsl8-2* (J,K). Differential-interference-contrast (G–I) or confocal-laser-scanning (J,K) microscopy. Bars: (B–D) 1 mm; (E) 0.25 mm; (G,H,J) 50 μm; (I,K) 25 μm.

*cals3-d* mutants formed narrow leaves whose vein networks deviated from those of WT in at least four respects: (1) fewer veins were formed; (2) closed loops were often replaced by open loops, i.e. loops that contact the midvein or other loops at only one of their two ends; (3) veins were often replaced by “vein fragments”, i.e. stretches of vascular elements that fail to contact other stretches of vascular elements at either one of their two ends, or by “vascular clusters”, i.e. isolated clusters of varied sizes and shapes, composed of improperly aligned and connected vascular elements; and (4) free vein ends often terminated in vascular clusters (Fig. 1C,D,F,H,I).

As *cals3-d*, *gsl8* mutants formed networks of fewer veins in which closed loops were often replaced by open loops; veins were often replaced by vein fragments or isolated vascular clusters; and free vein ends often terminated in vascular clusters (Fig. 1E,F,J,K). However, the vein fragments of strong *gsl8* mutants, such as *gsl8-2* and *gsl8 - chorus* (*gsl8-chor* hereafter), were shorter, and the clusters were rounder and larger than those of *cals3-d* and weak *gsl8* mutants such as *gsl8 - enlarged tetrad 2* (*gsl8-et2* hereafter) (Fig. 1E,F,J,K). Finally, the leaves of strong *gsl8* mutants were smaller than those of WT, *cals3-d*, and weak *gsl8* mutants; and in contrast to the entire leaf-margin of WT, *cals3-d*, and weak *gsl8* mutants, the leaf margin of strong *gsl8* mutants was lobed (Fig. 1E,F).

That defects in PD aperture regulation led to defects in vein formation, vascular element alignment and connection, and vein continuity and connection is consistent with the hypothesis that PD-mediated movement of an auxin signal controls vein patterning.

### PD Permeability Changes During Leaf Development

Because both near-constitutively wide and near-constitutively narrow PD aperture control vein patterning (Figure 1), we asked how PD permeability changed during leaf development. To address this question, we expressed an untargeted YFP, which diffuses through PDs (Imlau et al., 1999; Kim et al., 2005a; Kim et al., 2005b), by the *UAS* promoter, which is inactive in plants in the absence of a GAL4 driver (Fig. 2G). We activated YFP expression by tissue- and stage-specific GAL4/erGFP enhancer-trap (ET) drivers (Fig. 2F), and imaged erGFP expression and YFP signals in first leaves of the resulting ET>>erGFP/YFP plants 2.5 to 6 days after germination (DAG).

**Figure 2.**
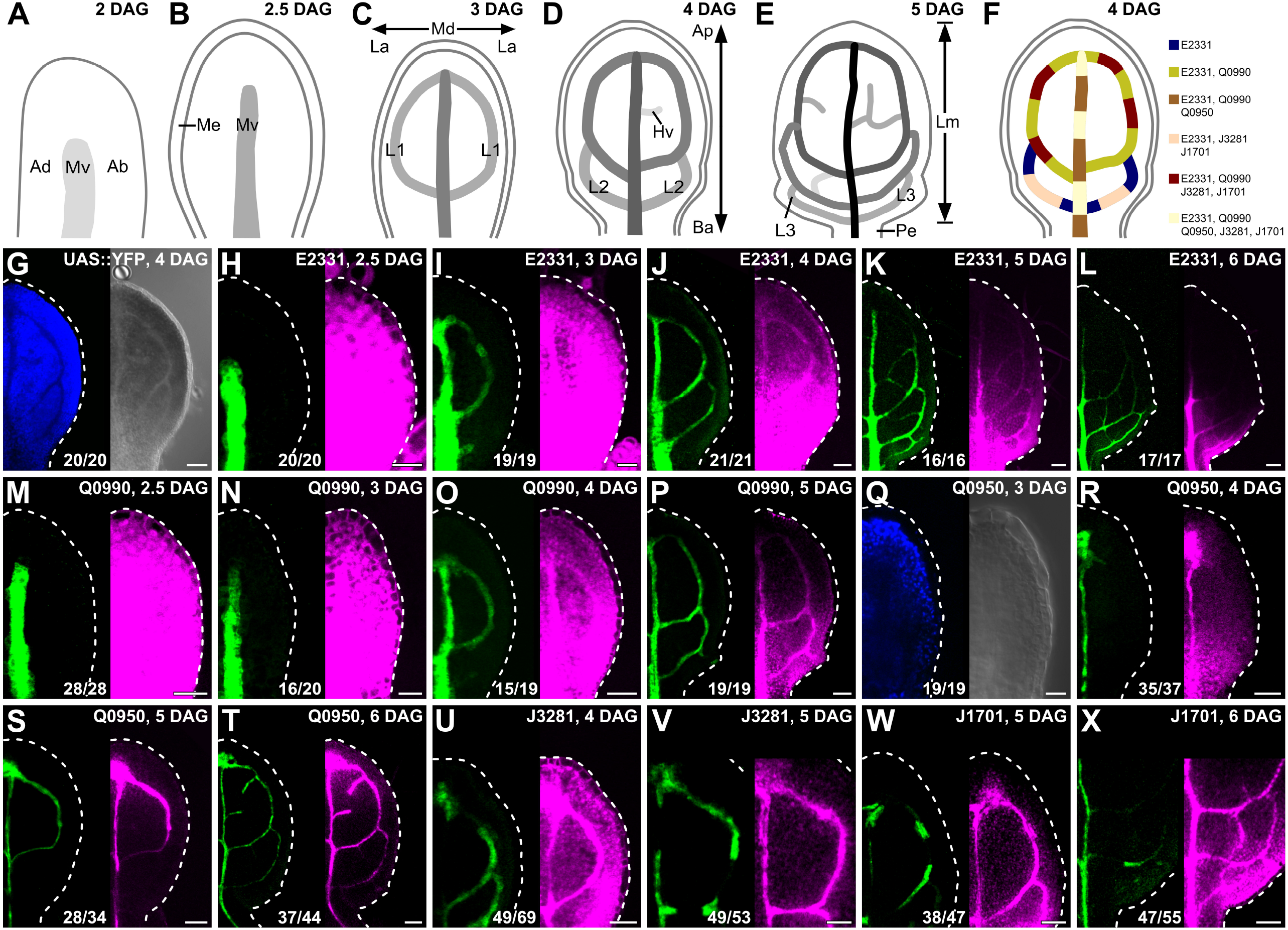
PD Aperture Changes During Leaf Development. (A–F) Top right: leaf age in days after germination (DAG). (A–E) Veins form sequentially during Arabidopsis leaf development: the formation of the midvein is followed by the formation of the first loops of veins (“first loops”); the formation of first loops is followed by that of second loops and minor veins; and the formation of second loops and minor veins is followed by that of third loops. Loops and minor veins form in a tip-to-base sequence during leaf development. Increasingly darker grays depict progressively later stages of vein development. Ab, abaxial; Ad, adaxial; Ap, apical; Ba, basal; Hv, minor vein; L1, L2, and L3: first, second, and third loop; La, lateral; Lm, lamina; Md, median; Me, marginal epidermis; Mv, midvein; Pe, petiole. (F) Expression map of tissue- and stage-specific GAL4/erGFP enhancer-trap (ET) drivers in developing leaves illustrates inferred overlap and exclusivity of expression. (G–X) Differential-interference-contrast (G, right; Q, right) or confocal-laser-scanning (all other panels) microscopy. First leaves (for simplicity, only half-leaves are shown). Blue, autofluorescence; green, GFP expression; magenta, YFP signals. Dashed white line delineates leaf outline. Top right: leaf age in DAG and transgene (G) or ET driver (all other panels). Bottom center: reproducibility index. Bars: (H,I,M,N,Q) 20 μm; (G,J,O,R,U) 40 μm; (K,P,S,V,W) 60 μm; (L,T,X) 80 μm.

In ET>>erGFP/YFP plants, expression of a nondiffusible endoplasmic-reticulum-localized GFP (erGFP) (Oparka et al., 1999) reports expression of GAL4 (Haseloff, 1999; Laplaze et al., 2005; Gardner et al., 2009), which activates expression of the diffusible YFP. Should the aperture of the PDs in the cells in which YFP expression is activated be narrower than the size of YFP, erGFP and YFP would be detected in the same cells. By contrast, should the aperture be wider than the size of YFP, YFP would be detected in additional cells.

The development of Arabidopsis leaves has been described previously (Pyke et al., 1991; Telfer and Poethig, 1994; Kinsman and Pyke, 1998; Candela et al., 1999; Donnelly et al., 1999; Mattsson et al., 1999; Kang and Dengler, 2002; Mattsson et al., 2003; Kang and Dengler, 2004; Scarpella et al., 2004). Briefly, at 2 DAG the first leaf is recognizable as a cylindrical primordium with a midvein at its center (Fig. 2A). By 2.5 DAG the primordium has expanded (Fig. 2B), and by 3 DAG the first loops of veins (“first loops”) have formed (Fig. 2C). By 4 DAG, a lamina and a petiole have become recognizable, and second loops have formed (Fig. 2D). By 5 DAG, lateral outgrowths have become recognizable in the lower quarter of the lamina; third loops have formed; and minor vein have formed in the upper three-quarters of the lamina (Fig. 2E). Finally, by 6 DAG minor vein formation has spread to the whole lamina.

In 2.5-DAG primordia of E2331>>erGFP/YFP, in which GAL4 expression is activated at early stages of vein development (Amalraj et al., 2020), YFP was detected throughout the 2.5-DAG primordium, even though erGFP was only expressed in the midvein (Fig. 2H). At 3 DAG, erGFP was only expressed in the midvein and first loops; nevertheless, YFP was still detected throughout the primordium, even though YFP signals were weaker at the primordium tip (Fig. 2I). At 4 DAG, erGFP was only expressed in the midvein and first and second loops (Fig. 2J). YFP signals were mainly restricted to the veins in the top half of the 4-DAG leaf but were detected throughout the bottom half of the 4-DAG leaf (Fig. 2J). At 5 DAG, erGFP was only expressed in the midvein; first, second, and third loops; and minor veins (Fig. 2K). YFP signals were mainly restricted to the veins in the upper three-quarters of the 5-DAG leaf but were detected throughout the lower quarter of the 5-DAG leaf (Fig. 2K). At 6 DAG, erGFP continued to be only expressed in the midvein; first, second, and third loops; and minor veins (Fig. 2L). YFP signals were mainly restricted to the veins in the whole 6-DAG leaf except for its lowermost part, where YFP was additionally detected in surrounding tissues (Fig. 2L).

YFP signals behaved during Q0990>>erGFP/YFP leaf development as they did during E2331>>erGFP/YFP leaf development (Fig. 2M–P; compare with Fig. 2H–L), even though GAL4 expression is activated at intermediate stages of vein development in Q0990, i.e. later than in E2331 (Sawchuk et al., 2007; Amalraj et al., 2020).

In 3-DAG primordia of Q0950>>erGFP/YFP, in which GAL4 is activated at late stages of vein development, i.e. later than in Q0990 (Sawchuk et al., 2007), neither erGFP nor YFP was expressed (Fig. 2Q), further suggesting that the *UAS* promoter is not active in plants in the absence of GAL4. At 4 DAG, erGFP was only expressed in the midvein (Fig. 2R). YFP signals were mainly restricted to the midvein in the top half of the 4-DAG leaf but were detected throughout the bottom half of the 4-DAG leaf (Fig. 2R). At 5 and 6 DAG, expression of both erGFP and YFP was restricted to the veins (Fig. 2S,T).

Consistent with previous observations (Kim et al., 2005a), our results suggest that PD permeability is high throughout the leaf at early stages of tissue development. PD permeability remains high in areas of the leaf where veins are still forming, but PD permeability between veins and surrounding nonvascular tissues lowers in areas of the leaf where veins are no longer forming. Eventually, veins become symplastically isolated from surrounding nonvascular tissues. To test whether vein cells become isolated also from one another, we imaged J3281>> and J1701>>erGFP/YFP, in which GAL4 is activated in vein segments (Sawchuk et al., 2007).

In 4- and 5-DAG J3281>>erGFP/YFP leaves, erGFP was only expressed in segments of midvein and first loops, but YFP was detected in the whole midvein and first loops (Fig. 2U,V). Likewise, in 5- and 6-DAG J1701>>erGFP/YFP leaves, erGFP was only expressed in segments of midvein and first and second loops, but YFP was detected in whole midvein and loops (Fig. 2W,X). These results suggest that vein cells are not symplastically isolated from one another even when they are isolated from surrounding nonvascular tissues.

### Auxin-Induced Vein Formation and PD Aperture Regulation

Auxin application induces vein formation (Sachs, 1989; Aloni, 2001; Scarpella et al., 2006; Sawchuk et al., 2007; Verna et al., 2019). Therefore, should the movement of an auxin signal that controls vein patterning be mediated by PDs, defects in PD aperture regulation would lead to defects in auxin-induced vein formation. To test this prediction, we applied the natural auxin indole-3-acetic acid (IAA) to one side of 3.5-DAG first leaves of E2331 and Q0990, and — because *cals3-2d* and *gsl8* leaves develop more slowly than WT leaves (see below) — of 4.5-DAG first leaves of E2331;*cals3-2d* and Q0990;*gsl8-2* (Fig. 3A,B,E,F). We then assessed erGFP-expression-labeled, IAA-induced vein formation 2.5 days later.

**Figure 3.**
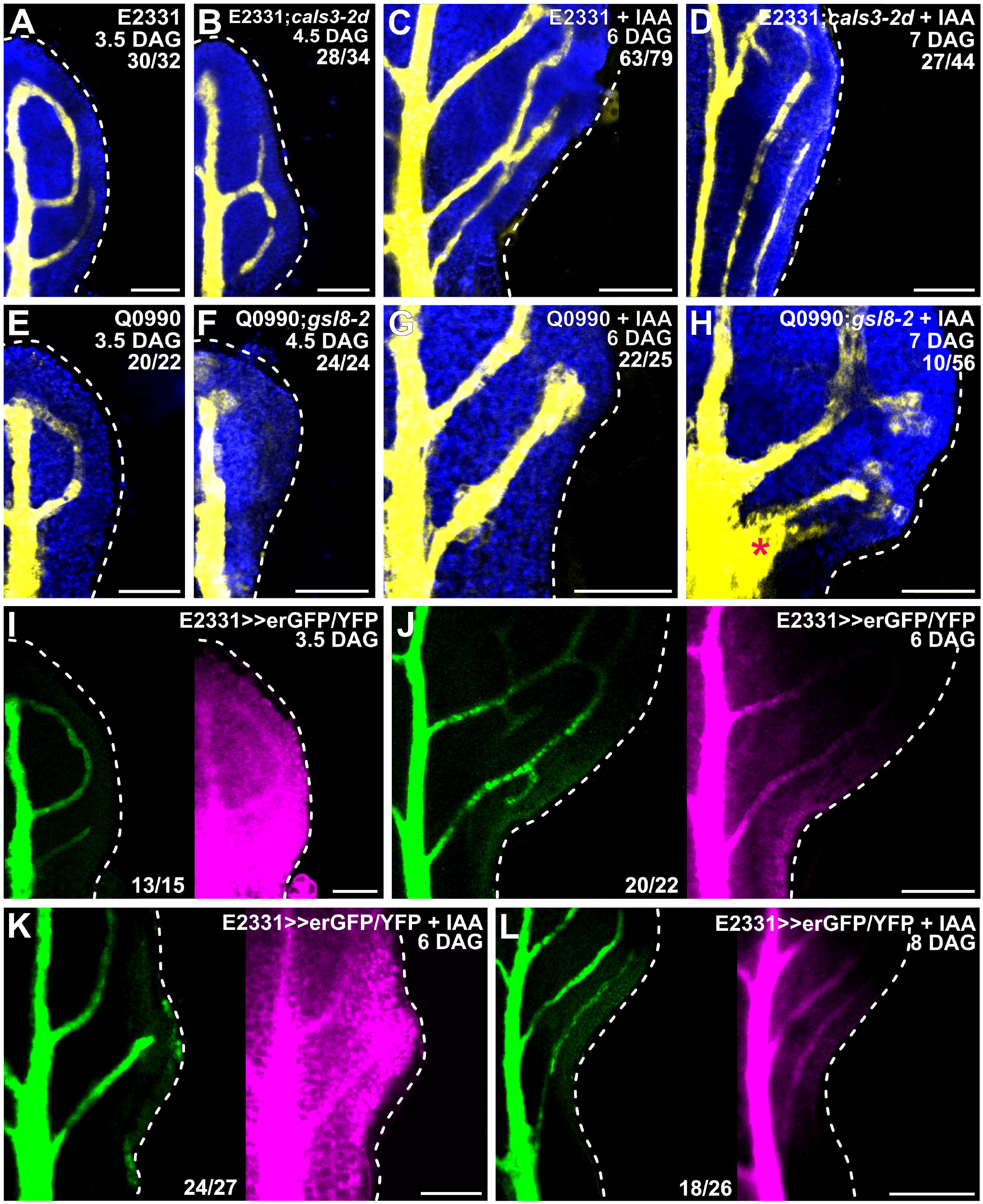
Auxin-Induced Vein Formation and PD Aperture Regulation. (A–L) Confocal laser scanning microscopy. First leaves (for simplicity, only half-leaves are shown). Blue, autofluorescence; yellow (A–H) or green (I–L), GFP expression; magenta, YFP signals. Dashed white line delineates leaf outline. Top right: leaf age in DAG, genotype, treatment, and — in A–H — reproducibility index. Bottom center (I-L): reproducibility index. Star in H indicates broad area of vascular differentiation connecting the midvein with the vein whose formation was induced by IAA application. Bars: (A,E,F,G–I,K) 80 μm; (B) 60 μm; (C,J,L) 120 μm; (D) 150 μm.

Consistent with previous reports (Sachs, 1989; Aloni, 2001; Scarpella et al., 2006; Sawchuk et al., 2007; Verna et al., 2019), IAA application induced the formation of veins in ∼80% (63/79) of E2331 leaves and ∼90% (22/25) of Q0990 leaves (Fig. 3C,G). Furthermore, in ∼55% (34/63) of the E2331 leaves and ∼65% (14/22) of the Q0990 leaves in which veins formed in response to IAA application, veins readily connected to the midvein (Fig. 3C,G). By contrast, IAA application induced vein formation in only ∼60% (27/44) of E2331;*cals3-2d* leaves and ∼20% (10/56) of Q0990;*gsl8-2* leaves (Fig. 3D,H). Moreover, only in ∼30% (8/27) of the E2331;*cals3-2d* leaves in which veins did form in response to IAA application did these veins connect to the midvein: in the remaining ∼70% of the responding leaves, the veins whose formation was induced by IAA application ran parallel to the midvein through the leaf petiole (Fig. 3D). Conversely, in 90% (9/10) of the Q0990;*gsl8-2* leaves in which IAA induced vein formation, not only did the veins whose formation was induced by IAA application connect to the midvein, but they did so by expanding into a broad vascular differentiation zone (Fig. 3H). Therefore, both near-constitutively wide and near-constitutively narrow PD aperture inhibit auxin-induced vein formation. Whenever the tissue escapes such an effect, near-constitutively narrow PD aperture inhibits connection of newly formed veins to pre-existing ones, and near-constitutively wide PD aperture accentuates that connection through excess vascular differentiation. These observations suggest that auxin-induced vein formation depends on regulated PD aperture, that restriction of auxin-induced vascular differentiation to limited cell files depends on narrow PD aperture, and that connection of veins whose formation is induced by auxin depends on wide PD aperture. Were that so, auxin application would impinge on the reduction in PD permeability between veins and surrounding nonvascular tissues that occurs during normal leaf development (Figure 2). To test this prediction, we applied IAA to one side of 3.5-DAG first leaves of E2331>>erGFP/YFP and assessed erGFP-expression-labeled, IAA-induced vein formation and YFP-signal-inferred PD permeability 2.5 and 4.5 days later (i.e. 6 and 8 DAG, respectively).

Consistent with what shown above (Fig. 2I,J; Fig. 3A), at 3.5 DAG erGFP was only expressed in the midvein and in first and second loops (Fig. 3I). In the top half of 3.5-DAG leaves, YFP signals were weaker in nonvascular tissues than in veins but were uniformly strong in the bottom half of the leaves (Fig. 3I). As shown above (Fig. 2L), at 6 DAG erGFP expression was restricted to the veins (Fig. 3J,K). In 6-DAG leaves to which IAA had not been applied, YFP signals were mainly restricted to the veins in the whole leaf except for its lowermost part, where YFP was additionally detected in surrounding tissues (Fig. 3J). By contrast, YFP signals were detected throughout the bottom half of the 6-DAG leaves to which IAA had been applied (Fig. 3K). By 8 DAG, however, both erGFP and YFP signals had become mainly restricted to the veins also in the leaves to which IAA had been applied (Fig. 3L).

In conclusion, our results suggest that auxin application delays the reduction in PD permeability between veins and surrounding nonvascular tissues that occurs during normal leaf development, and that auxin-induced vein formation and connection depends on the ability to regulate PD aperture. Such conclusions are consistent with the hypothesis that the movement of an auxin signal that controls vein patterning is mediated by PDs.

### Auxin-Transport-Dependent Vein Patterning and Regulated PD Aperture

Should the auxin-transporter-unmediated movement of an auxin signal that controls vein patterning be mediated by PDs, defects in PD aperture regulation would enhance vein patterning defects induced by auxin transport inhibition. To test this prediction, we grew E2331, E2331;*cals3-2d*, and E2331;*gsl8-2* in the presence or absence of the auxin transport inhibitor N-1-naphthylphthalamic acid (NPA) (Morgan and Söding, 1958), which induces vein patterning defects that phenocopy the loss of that auxin transport pathway that is relevant for vein patterning (Verna et al., 2019). We then imaged erGFP-expression-labeled vein networks 4.5 DAG in E2331 and — because *cals3-2d* and *gsl8* leaves develop more slowly than WT leaves (see below) — 5.5 and 7 DAG in E2331;*cals3-2d* and E2331;*gsl8-2*, respectively.

Consistent with what shown above (Fig. 2J,K), in the absence of NPA vein networks were composed of midvein, first and second loops, and minor veins in E2331, and of midvein, loops — whether open or closed — vein fragments, and vascular clusters in E2331;*cals3-2d* and E2331;*gsl8-2* (Fig. 4A–C).

**Figure 4.**
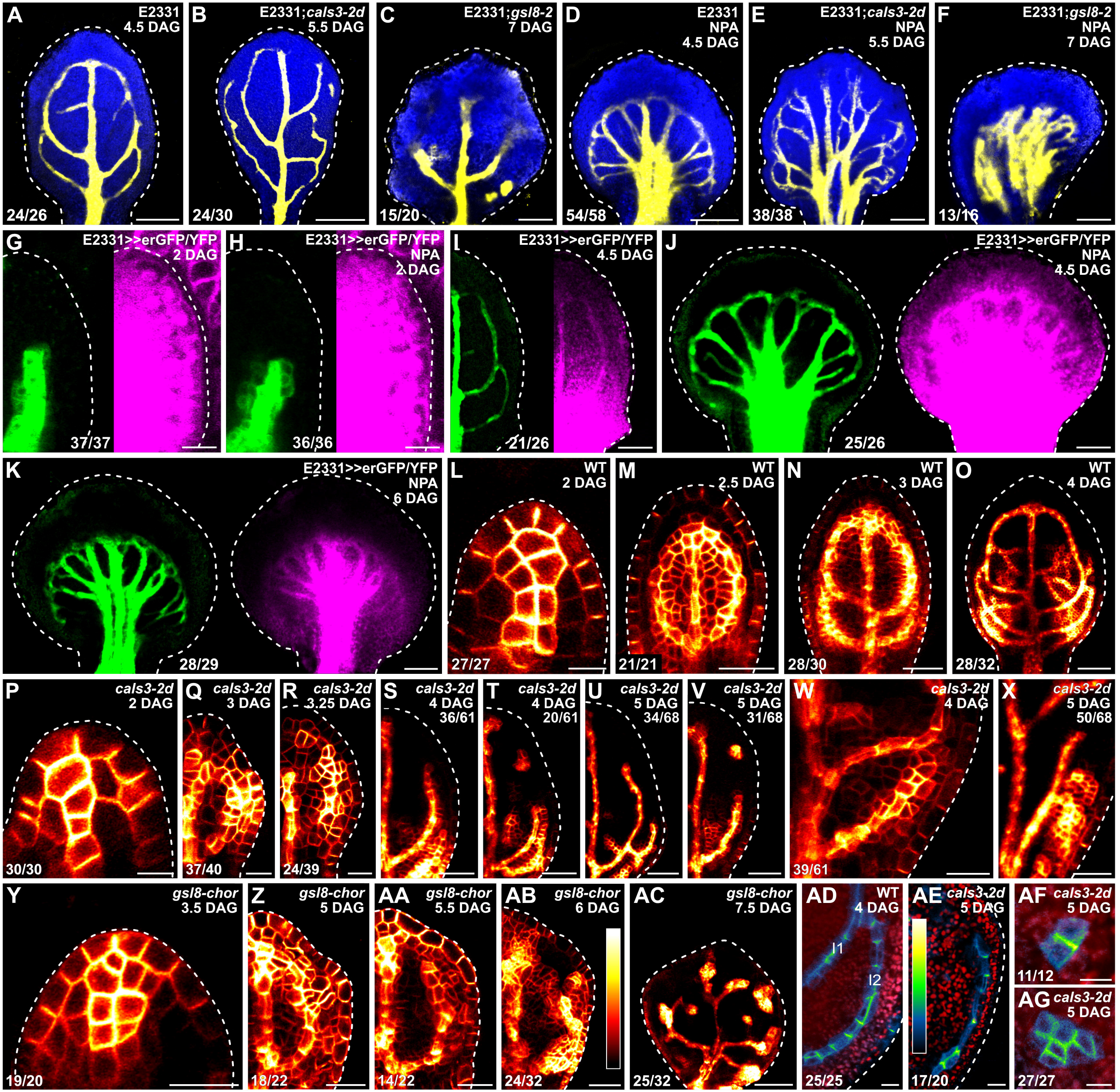
Auxin-Transport-Dependent Vein Patterning and Regulated PD Aperture. (A–AG) Confocal laser scanning microscopy. First leaves (for simplicity, only half-leaves are shown in G–I, Q–V, and Z–AB). Blue (A–F) or red (AD–AG), autofluorescence; yellow (A–F) or green (G–K), GFP expression; magenta, YFP signals. Dashed white line delineates leaf outline. (G,H,L,P,Y) Side view, adaxial side to the left. (L–AC) PIN1::PIN1:YFP expression; look-up table (ramp in AB) visualizes expression levels. (AD–AG) PIN1::PIN1:GFP (AD,AF,AG) or PIN1::PIN1:YFP (AE) expression; look-up table (ramp in AE) visualizes expression levels. Top right: leaf age in DAG, genotype, and treatment (25 μM NPA), and — in S–V,X — reproducibility index. Bottom left (A–F,L– R,W,Y–AG) or center (G–K): reproducibility index. Bars: (A–F,I–K) 120 μm; (G,H,Q,R,W,X,Y) 20 μm; (L,P,AD–AG) 10 μm; (M,N,S,T,Z,AA,AB) 40 μm; (O,U,V,AC) 60 μm.

Consistent with previous reports (Mattsson et al., 1999; Sieburth, 1999; Verna et al., 2019), NPA reproducibly induced characteristic vein-pattern defects in E2331 leaves: (1) the vein network comprised more lateral-veins; (2) lateral veins failed to join the midvein in the middle of the leaf and instead ran parallel to one another to form a wide midvein; and (3) lateral veins joined distal veins in a marginal vein that closely paralleled the leaf margin and gave a smooth outline to the vein network (Fig. 4D).

As in E2331, in E2331;*cals3-2d* NPA induced the formation of more lateral-veins that failed to join the midvein in the middle of the leaf and instead ran parallel to one another to form a wide midvein (Fig. 4E). However, unlike in NPA-grown E2331, in NPA-grown E2331;*cals3-2d* lateral veins often failed to join distal veins in a marginal vein and instead ended freely in the lamina near the leaf margin (Fig. 4E).

As in both E2331 and E2331;*cals3-2d*, in E2331;*gsl8-2* NPA induced the formation of more veins, but these veins ran parallel to one another to give rise to a midvein that spanned almost the entire width of the leaf (Fig. 4F). And as in NPA-grown E2331;*cals3-2d* — but unlike in NPA-grown E2331 — in NPA-grown E2331;*gsl8-2* only very rarely did veins join one another in a marginal vein; instead, they most often ended freely in the lamina near the leaf tip (Fig. 4F).

In conclusion, both near-constitutively wide and near-constitutively narrow PD aperture enhance vein patterning defects induced by auxin transport inhibition, a conclusion that is consistent with the hypothesis that the auxin-transporter-unmediated movement of an auxin signal that controls vein patterning is mediated by PDs. Moreover, because auxin transport inhibition promotes vein connection (Verna et al., 2015), that NPA was unable to induce vein connection in E2331;*cals3-2d* and E2331;*gsl8-2* suggests that the promoting effect of auxin transport inhibition on vein connection is mediated by regulated PD permeability. Were that so, auxin transport inhibition would impinge on the reduction of PD permeability that occurs between veins and surrounding nonvascular tissues during normal leaf development. To test this prediction, we grew E2331>>erGFP/YFP in the presence or absence of NPA, and assessed erGFP-expression-labeled vein network formation and YFP-signal-inferred PD aperture in first leaves 2, 4.5, and 6 DAG.

Consistent with what shown above (Fig. 2H), at 2 DAG erGFP was only expressed in the midvein of both NPA- and normally grown E2331>>erGFP/YFP — though the erGFP expression domain was broader in NPA-grown than in normally grown primordia (Fig. 4G,H). Likewise, in both NPA- and normally grown E2331>>erGFP/YFP YFP was detected throughout the 2-DAG primordia (Fig. 4G,H).

Also consistent with what shown above (Fig. 2J,K), at 4.5 DAG erGFP was only expressed in the midvein, first and second loops, and minor veins of normally grown E2331>>erGFP/YFP (Fig. 4I). YFP signals were mainly restricted to the veins in the top half of normally grown 4.5-DAG E2331>>erGFP/YFP leaves but were detected throughout the bottom half of the leaves (Fig. 4I). Also in NPA-grown 4.5-DAG E2331>>erGFP/YFP leaves, erGFP expression was restricted to the veins; however, YFP was detected throughout NPA-grown 4.5-DAG E2331>>erGFP/YFP leaves — though YFP signals were weaker along the margin in the top half of the leaves (Fig. 4J). Nevertheless, by 6 DAG both erGFP and YFP signals had become mainly restricted to the veins also in NPA-grown E2331>>erGFP/YFP (Fig. 4K).

We conclude that auxin transport inhibition delays the reduction in PD permeability between veins and surrounding nonvascular tissues that occurs during normal leaf development, and that such delay mediates the promoting effect of auxin transport inhibition on vein connection.

We next asked whether regulated PD aperture controlled polar auxin transport during leaf development. Because *PIN1* is the only auxin-transporter-encoding gene in Arabidopsis with nonredundant functions in vein patterning (Sawchuk et al., 2013), to address that question we imaged expression domains and cellular localization of expression of PIN1::PIN1:YFP (PIN1:YFP fusion protein expressed by the *PIN1* promoter (Xu et al., 2006)) or PIN1::PIN1:GFP (Benkova et al., 2003) during first-leaf development in WT, *cals3-2d*, and *gsl8-chor*.

Consistent with previous reports (Benkova et al., 2003; Reinhardt et al., 2003; Heisler et al., 2005; Scarpella et al., 2006; Wenzel et al., 2007; Bayer et al., 2009; Sawchuk et al., 2013; Marcos and Berleth, 2014; Verna et al., 2015; Verna et al., 2019), in WT PIN1::PIN1:YFP was expressed in all the cells at early stages of tissue development, and inner tissue expression was stronger in developing veins (Fig. 4L–O). Over time, epidermal expression became restricted to the basalmost cells, and inner tissue expression became restricted to developing veins (Fig. 4L–O).

Also in *cals3-2d* and *gsl8-chor*, PIN1::PIN1:YFP was expressed in all the cells at early stages of tissue development, and inner tissue expression was stronger in developing veins (Fig. 4P–AC). Furthermore, as in WT, in both *cals3-2d* and *gsl8-chor* PIN1::PIN1:YFP expression domains associated with loop formation were initially connected on both ends to pre-existing expression domains, and PIN1::PIN1:YFP was evenly expressed along those looped domains (Fig. 4Q,Z). However, in *casl3-2d* and *gsl8-chor* PIN1::PIN1:YFP expression along looped domains soon became heterogeneous, with domain segments with stronger expression separated by segments with weaker expression (Fig. 4R,W,AA,AB). Such heterogeneity in PIN1::PIN1:YFP expression at early stages of loop formation was associated with open or fragmented looped domains of PIN1::PIN1:YFP expression at later stages (Fig. 4S–V,X,AC). Finally, equivalent stages of vein development occurred at later time points in *casl3-2d* and *gsl8-chor* than in WT (e.g., compare Fig. 4Q,Z with Fig. 4M, Fig. 4S,T,W with Fig. 4N, and Fig. 4U,V,X,AC with Fig. 4O).

Consistent with previous reports (Scarpella et al., 2006; Wenzel et al., 2007; Bayer et al., 2009; Sawchuk et al., 2013; Marcos and Berleth, 2014; Verna et al., 2015; Verna et al., 2019), in cells at late stages of second loop development in WT leaves, by which time PIN1::PIN1:GFP expression had become restricted to the cells of the developing loop, PIN1::PIN1:GFP expression was localized to the side of the plasma membrane facing the veins to which the second loop was connected (Fig. 4AD).

By contrast, in cells at late stages of development of vein fragments and isolated vascular clusters in *cals3-2d*, by which time expression domains of PIN1::PIN1:GFP or PIN1::PIN1:YFP (hereafter collectively referred to as PIN1::PIN1:FP) had become disconnected from other veins on both ends, PIN1::PIN1::FP expression was localized to any of the plasma membrane sides facing a contiguous PIN1::PIN1:FP-expressing cell (Fig. 4AE-AG).

In conclusion, our results suggest that vein patterning is controlled by the mutually coordinated action of polar auxin transport and PD-mediated movement of an auxin signal.

### Auxin-Signaling-Dependent Vein Patterning and Regulated PD Aperture

The auxin-transporter-unmediated movement of an auxin signal that controls vein patterning is, at least in part, mediated by auxin signaling (Verna et al., 2019). Should the residual, auxin-transporter- and auxin-signaling-unmediated movement of an auxin signal that controls vein patterning be mediated by PDs, defects in PD aperture regulation would enhance vein patterning defects induced by auxin signaling inhibition — just as defects in PD aperture enhance vein patterning defects induced by auxin transport inhibition (Fig. 4A–F). To test the prediction that defects in PD aperture will enhance vein patterning defects induced by auxin signaling inhibition, we grew E2331, E2331;*cals3-2d*, and E2331;*gsl8-2* in the presence or absence of the auxin signaling inhibitor phenylboronic acid (PBA) (Matthes and Torres-Ruiz, 2016), which induces vein patterning defects that phenocopy the loss of that auxin signaling pathway that is relevant for vein patterning (Verna et al., 2019). We then imaged erGFP-expression-labeled vein networks 4.5 DAG in E2331, and — because *cals3-2d* and *gsl8* leaves develop more slowly than WT leaves (Fig. 4M–O,Q,S–X,Z,AC) — 5.5 and 6.5 DAG in E2331;*cals3-2d* and E2331;*gsl8-2*, respectively.

As shown above (Fig. 4A–C), in the absence of PBA vein networks were composed of midvein, first and second loops, and minor veins in E2331, and of midvein, loops — whether open or closed — vein fragments, and vascular clusters in E2331;*cals3-2d* and E2331;*gsl8-2* (Fig. 5A,D,G).

**Figure 5.**
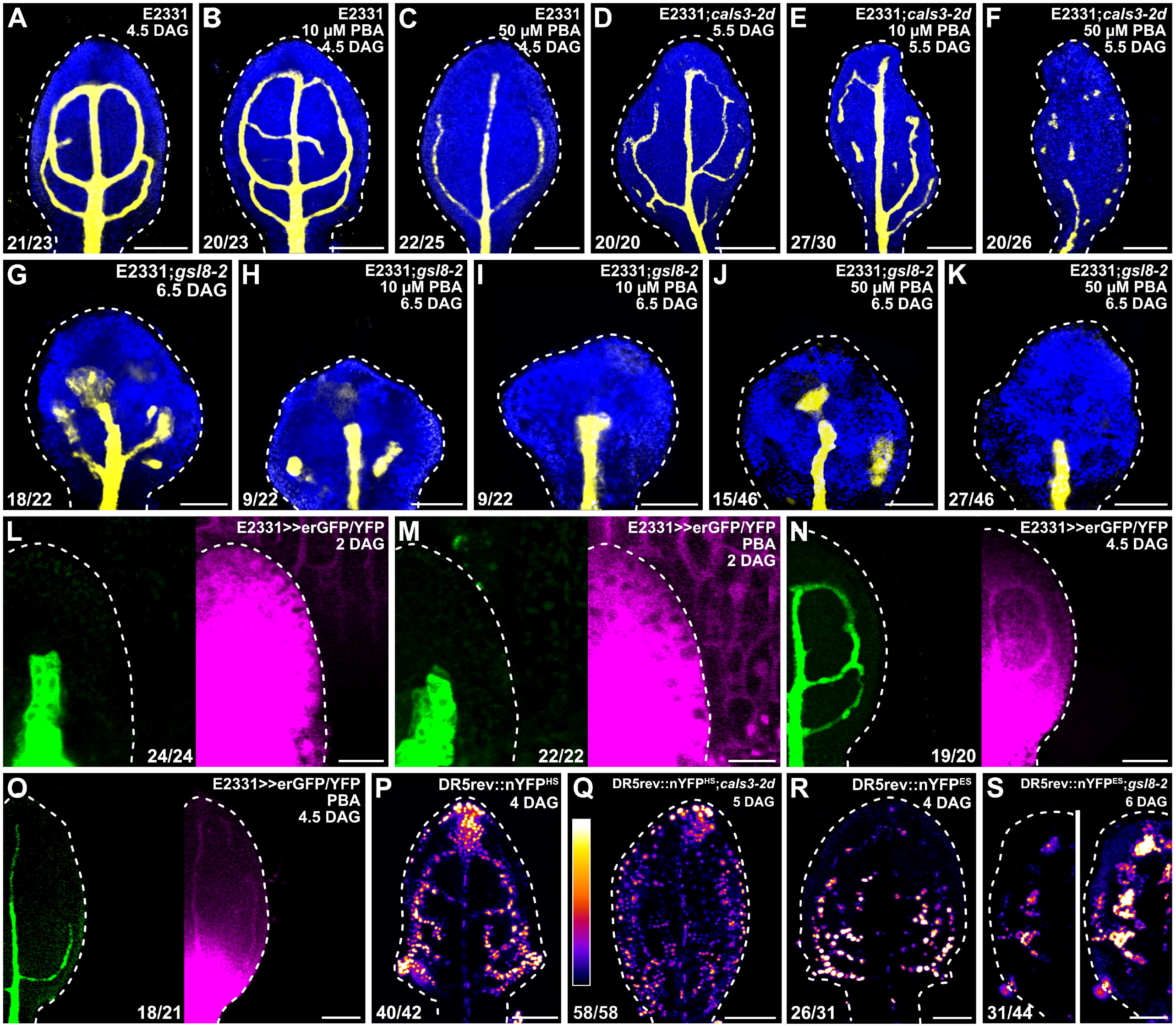
Auxin-Signaling-Dependent Vein Patterning and Regulated PD Aperture. (A–S) Confocal laser scanning microscopy. First leaves (for simplicity, only half-leaves are shown in L–O,S). Blue, autofluorescence; yellow (A– K) or green (L–O), GFP expression; magenta, YFP signals. (P–S) DR5rev::nYFP^HS^ (P,Q) or DR5rev::nYFP^ES^ (R,S) expression; look-up table (ramp in Q) visualizes expression levels. Dashed white line delineates leaf outline. Top right: leaf age in DAG, genotype, and treatment (10 or 50 μM PBA). Bottom left (A–K,P–S) or center (L–O): reproducibility index. Images in P and R were acquired by matching signal intensity to detector’s input range (∼1% saturated pixels). Images in P and Q were acquired at identical settings and show weaker and broader DR5rev::nYFP^HS^ expression in *cals3-2d*. Images in R and S (left) were acquired at identical settings and show weaker DR5rev::nYFP^ES^ expression in *gsl8-2*. Image in S (right) was acquired by matching signal intensity to detector’s input range (∼1% saturated pixels), and show broader DR5rev::nYFP^ES^ expression in *gsl8-2*. Bars: (A–K) 120 μm; (L,M) 20 μm; (N–S) 80 μm.

Ten μM PBA failed to induce vein network defects in E2331 but led to the formation of fewer veins and opening or fragmentation of all the loops in E2331;*cals3-2d* and E2331;*gsl8-2* (Fig. 5B,E,H,I). Formation of fewer veins and opening of all the loops — though not their fragmentation — were induced in E2331 by 50 μM PBA (Fig. 5C). At that concentration of PBA, the vascular systems of E2331;*cals3-2d* and E2331;*gsl8-2* were mainly composed of very few, scattered vascular clusters (Fig. 6F,J,K).

**Figure 6.**
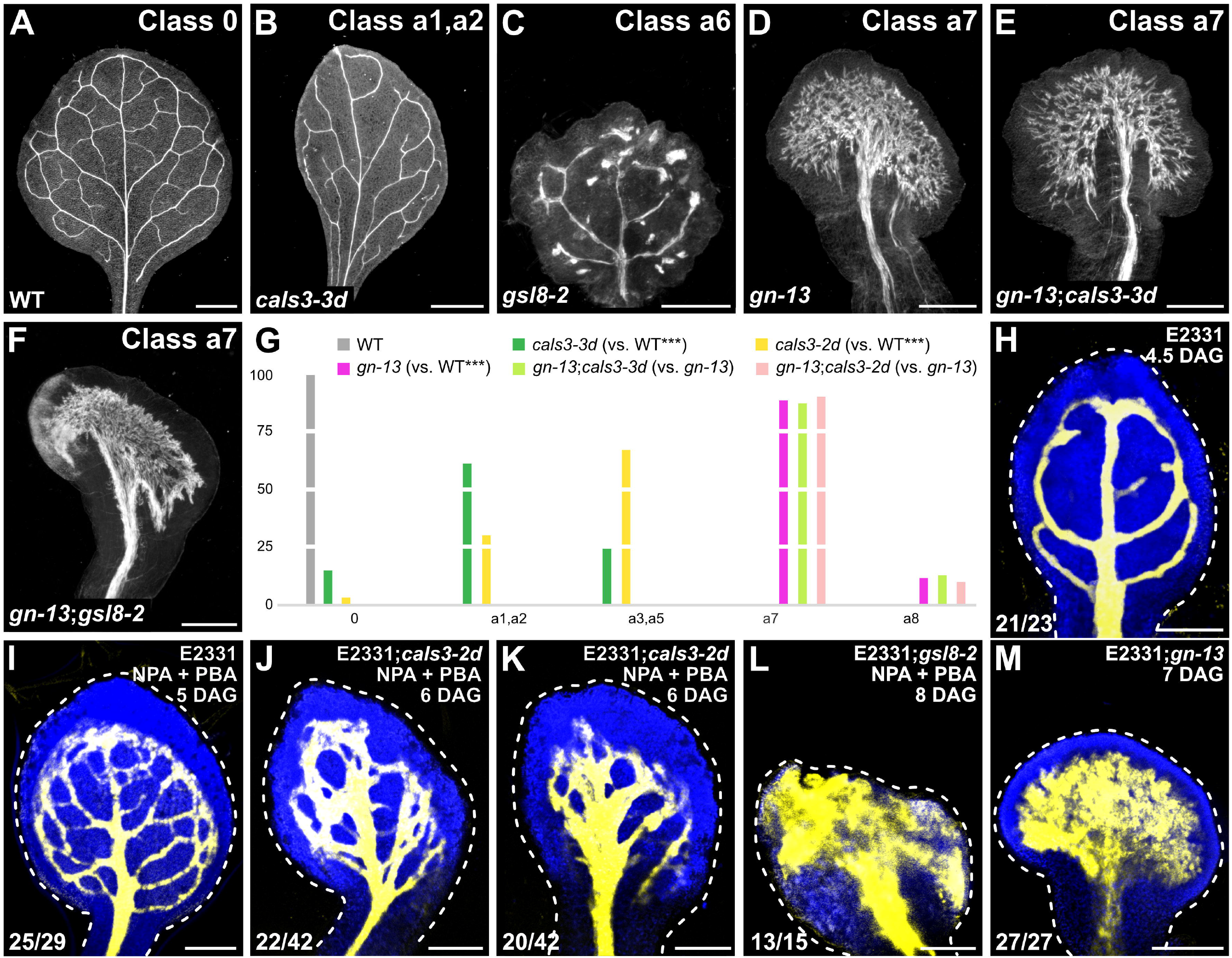
Control of PD-Mediated Vein Patterning by *GNOM*. (A–F) Dark-field illumination of mature first leaves illustrating phenotype classes (top right) and genotypes (bottom left). Classes 0, a1, a2, and a6 defined in Figure 1. Class a7: wide midvein and shapeless vascular cluster (D–F). The *gn* phenotype segregated in approximately one-quarter (559/2,353) of the progeny of plants heterozygous for both *gn-13* and *gsl8-2* — no different from the frequency expected by the Pearson’s chi-squared (χ2) goodness-of-fit test for the hypothesis that the phenotype of *gn-13* is epistatic to that of *gsl8-2*. We genotyped 68 *gn*-looking seedlings and found that four were *gn-13*;*gsl8-2* double homozygous mutants. (G) Percentages of leaves in phenotype classes. Classes 0, a1–a3, a5, and a6 defined in Figure 1. Class a8, shapeless vascular cluster (not shown). Difference between *cals3-3d* and WT, between *cals3-2d* and WT, and between *gn-13* and WT was significant at *P*<0.001 (***) by Kruskal-Wallis and Mann-Whitney test with Bonferroni correction. Sample population sizes: WT, 30; *cals3-3d*, 62; *cals3-2d*, 67; *gn-13*, 89; *gn-13*;*cals3-3d*, 100; *gn-13*;*cals3-2d*, 52. (H–M) Confocal laser scanning microscopy. First leaves. Blue, autofluorescence; yellow, GFP expression. Dashed white line delineates leaf outline. Top right: leaf age in DAG, genotype, and treatment (25 μM NPA + 10 μM PBA). Bottom left: reproducibility index. Bars: (A,B) 1 mm; (C,D,F) 0.5 mm; (E) 0.25 mm; (H) 100 μm; (I,L,M) 150 μm; (J,K) 100 μm.

These observations suggest that defects in PD aperture enhance vein patterning defects induced by auxin signaling inhibition, a conclusion that is consistent with the hypothesis that the residual, auxin-transporter- and auxin-signaling-unmediated movement of an auxin signal that controls vein patterning is mediated by PDs. Moreover, these observations suggest that the vein patterning defects induced by auxin signaling inhibition may, at least in part, be mediated by regulated PD permeability. Were that so, auxin signaling inhibition would impinge on the reduction of PD permeability that occurs between veins and surrounding nonvascular tissues during normal leaf development. To test this prediction, we grew E2331>>erGFP/YFP in the presence or absence of PBA, and assessed erGFP-expression-labeled vein network formation and YFP-signal-inferred PD aperture in first leaves 2 and 4.5 DAG.

As shown above (Fig. 4G), at 2 DAG erGFP was only expressed in the midvein of both PBA- and normally grown E2331>>erGFP/YFP (Fig. 5L,M). Likewise, in both PBA- and normally grown E2331>>erGFP/YFP YFP was detected throughout the 2-DAG primordia (Fig. 5L,M).

As also shown above (Fig. 4I), at 4.5 DAG erGFP was only expressed in the midvein, first and second loops, and minor veins of normally grown E2331>>erGFP/YFP (Fig. 5N). YFP signals were mainly restricted to the veins in the top half of normally grown 4.5-DAG E2331>>erGFP/YFP leaves but were detected throughout the bottom half of the leaves (Fig. 5N). Also in PBA-grown 4.5-DAG E2331>>erGFP/YFP leaves, erGFP expression was restricted to the veins; however, YFP signals were already restricted to the veins in the top three-quarters of PBA-grown 4.5-DAG E2331>>erGFP/YFP leaves (Fig. 5O), suggesting that auxin signaling inhibition leads to premature reduction in PD permeability.

We conclude that auxin signaling inhibition prematurely reduces PD permeability between veins and surrounding nonvascular tissues, and that such premature reduction mediates, at least in part, the effects of auxin signaling inhibition on vein patterning.

We next asked whether regulated PD aperture in turn controlled response to auxin signals in developing leaves. To address this question, we imaged expression of the auxin response reporter DR5rev::nYFP (Heisler et al., 2005; Sawchuk et al., 2013) (Table S1) 4 DAG in WT and — because *cals3-2d* and *gsl8* leaves develop more slowly than WT leaves (Fig. 4M–O,Q,S–X,Z,AC) — 5 and 6 DAG in *cals3-2d* and *gsl8-2*, respectively.

As previously shown (Sawchuk et al., 2013; Verna et al., 2015; Verna et al., 2019), in WT strong DR5rev::nYFP expression was mainly associated with developing veins (Fig. 5P,R). By contrast, DR5rev::nYFP expression was weaker and expression domains were broader in *cals3-2d* and *gsl8-2* (Fig. 5Q,S).

In conclusion, our results suggest that vein patterning is controlled by the mutually coordinated action of auxin signaling and PD-mediated movement of an auxin signal.

### Control of PD-Mediated Vein Patterning by GN

Vein patterning is controlled by the mutually coordinated action of auxin signaling, polar auxin transport, and PD-mediated movement of an auxin signal (Figures 1–5). Vein patterning activities of both auxin signaling and polar auxin transport depend on *GN* function (Verna et al., 2019). We asked whether *GN* also controlled PD-mediated vein patterning. To address this question, we compared the phenotypes of mature first leaves of the *gn-13*;*cals3-2d*, *gn-13*;*cals3-3d*, and *gn-13*;*gsl8-2* double mutants with those of their respective single mutants.

The phenotypes of *gn-13*;*cals3-2d*, *gn-13*;*cals3-3d*, and *gn-13*;*gsl8-2* were no different from those of *gn-13* (Fig. 6A–G), suggesting that the effects of the *gn-13* mutation on vein patterning are epistatic to those of the *cals3-d* or *gsl8-2* mutation. Therefore, *GN* controls PD-mediated vein patterning just as it controls auxin-transport- and auxin-signaling-dependent vein patterning.

Vein pattern defects of intermediate alleles of *gn* are phenocopied by growth in the presence of both NPA and PBA (Verna et al., 2019) (Fig. 6I). Because *GN* controls PD-mediated vein patterning in addition to auxin-transport- and auxin-signaling-dependent vein patterning, we asked whether defects in PD aperture shifted the defects induced by NPA and PBA toward more severe classes of the *gn* vein patterning phenotype. To address this question, we grew E2331, E2331;*cals3-2d*, and E2331;*gsl8-2* in the presence or absence of NPA and PBA. We then imaged erGFP-expression-labeled vein networks 5 DAG in E2331 and — because *cals3-2d* and *gsl8* leaves develop more slowly than WT leaves (Fig. 4M–O,Q,S– X,Z,AC) — 6 and 8 DAG in E2331;*cals3-2d* and E2331;*gsl8-2*, respectively.

Growth of *cals3-2d* in the presence of both NPA and PBA led to vein pattern defects similar to those of the strong *gn-van7* allele (Koizumi et al., 2000; Verna et al., 2019) (Fig. 6J,K), and growth of *gsl8-2* in the presence of both NPA and PBA even phenocopied vein pattern defects of the null *gn-13* allele (Fig. 6L,M).

We conclude that vein patterning is controlled by the *GN*-dependent, coordinated action of auxin signaling, polar auxin transport, and PD-mediated movement of an auxin signal.

## DISCUSSION

Unlike the tissue networks of animals, the vein networks of plant leaves form in the absence of cell migration and direct cell–cell interaction. Therefore, leaf vein networks are patterned by a mechanism unrelated to that which patterns animal tissue networks.

Here we show that leaf veins are patterned by the coordinated action of three *GN*-dependent pathways: auxin signaling, polar auxin transport, and movement of an auxin signal through PDs (Fig. 7F). Because in the most severe cases inhibition of those three pathways prevents patterning of vascular cells into veins, we have identified all the main pathways that control vein patterning.

**Figure 7.**
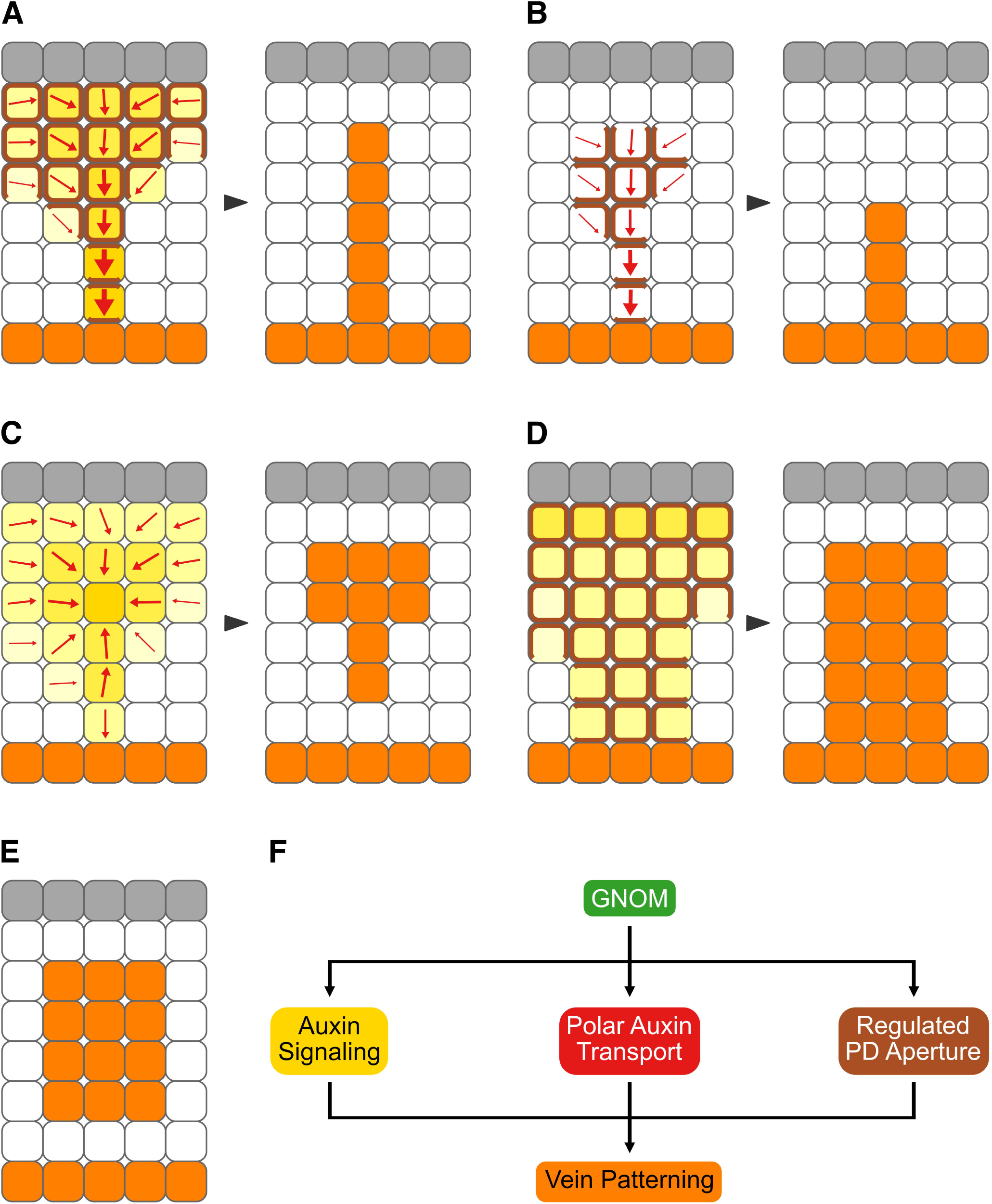
Summary and Interpretation. (A–F) Gray: epidermis, whose role in vein patterning — if any — remains unclear (Govindaraju et al., 2020). Increasingly darker yellow: progressively stronger auxin signaling. Increasingly thicker arrows: progressively more polarized auxin transport. Brown: PD-mediated cell– cell connection. Orange: veins. Arrowheads temporally connect vein patterning stages with mature vein patterns. (A) In WT, veins are patterned by gradual restriction of auxin signaling domains (Wenzel et al., 2007; Donner et al., 2009; Krogan et al., 2012; Esteve-Bruna et al., 2013; Marcos and Berleth, 2014; Krishna et al., 2021), gradual restriction of auxin transport domains and polarization of auxin transport paths (Scarpella et al., 2006; Wenzel et al., 2007; Bayer et al., 2009; Krogan et al., 2012; Esteve-Bruna et al., 2013; Sawchuk et al., 2013; Marcos and Berleth, 2014; Verna et al., 2015; Verna et al., 2019), and gradual reduction of PD permeability between incipient veins and surrounding nonvascular tissues. (B) Inhibition of auxin signaling leads to narrower domains of auxin transport (Wenzel et al., 2007; Esteve-Bruna et al., 2013; Verna et al., 2019) and promotes reduction of PD permeability between incipient veins and surrounding nonvascular tissues. (C) Defects in the ability to regulate PD aperture lead to weaker and broader domains of auxin signaling, fragmentation of auxin transport domains, and abnormal polarization of auxin transport paths. (D) Inhibition of auxin transport leads to weaker and broader domains of auxin signaling (Mattsson et al., 2003; Esteve-Bruna et al., 2013; Verna et al., 2015; Verna et al., 2019) and delays reduction of PD permeability between incipient veins and surrounding nonvascular tissues. (E) Loss of *GN* function or simultaneous inhibition of auxin signaling, polar auxin transport, and ability to regulate PD aperture leads to clusters of vascular cells. (F) Veins are patterned by the coordinated activities of three *GN*-dependent pathways: auxin signaling, polar auxin transport, and regulated PD aperture.

### Regulation of PD Permeability During Leaf Development

At early stages of leaf tissue development — stages at which veins are forming — PD permeability is high throughout the leaf (Fig. 7A). As leaf tissues develop, PD permeability between veins and surrounding nonvascular tissues becomes gradually lower, but PD permeability between vein cells remains high. These results suggest that at early stages of leaf tissue development all cells are symplastically connected; as veins develop, vein cells remain symplastically connected but become isolated from the surrounding nonvascular tissues.

The changes in PD permeability that occur during leaf development resemble those observed during the development of embryos (Kim et al., 2005a; Kim et al., 2005b; Stadler et al., 2005; Godel-Jedrychowska et al., 2020), lateral roots (Benitez-Alfonso et al., 2013; Sager et al., 2020), and stomata (Willmer and Sexton, 1979; Wille and Lucas, 1984; Palevitz and Hepler, 1985). By contrast, the changes in PD permeability that occur during leaf development are unlikely to be related to those observed during the transition of leaf tissues from sink to source of photosynthates as this transition begins when new veins are no longer forming and all existing veins have completely differentiated (e.g., (Roberts et al., 1997; Imlau et al., 1999; Oparka et al., 1999; Roberts et al., 2001; Wright et al., 2003)).

Consistent with the observation that vein formation is associated first with high and then with low PD permeability between veins and surrounding nonvascular tissues, defects in the ability to regulate PD aperture — whether leading to near-constitutively wide or narrow PD aperture — lead to similar vein patterning defects: fewer veins form, and those that do form become disconnected and discontinuous. In the most extreme cases, randomly oriented vascular elements differentiate in clusters, a phenotype that so far had only been observed in *gn* mutants or in plants impaired in both auxin signaling and polar auxin transport (Mayer et al., 1993; Koizumi et al., 2000; Geldner et al., 2004; Verna et al., 2019).

How symplastic connection between vein cells and their isolation from surrounding nonvascular tissues is brought about during vein development remains to be understood. One possibility is that, as in cells of the embryonic hypocotyl (Kim et al., 2005b), there are more PDs along the transverse walls of vein cells than along their longitudinal walls — perhaps because no new PDs form in the longitudinal walls of vein cells during their elongation, as it happens in elongating root cells (Gunning, 1978; Zhu et al., 1998). One other possibility is that, as it happens to elongating root cells (Seagull, 1983), during vein cell elongation simple PDs coalesce into branched PDs, which have narrower aperture (Oparka et al., 1999). Consistent with this possibility, there are more branched PDs along the longitudinal walls than along the transverse walls of epidermal cells underlying the midvein (Gao et al., 2020). Yet another possibility is that the aperture of PDs along the longitudinal walls of vein cells is narrower than that of PDs along their transverse walls — perhaps because the aperture of PDs along longitudinal walls is closed by the same turgor pressure that drives cell elongation (Oparka and Prior, 1992; Ruan et al., 2001; Park et al., 2019) or because more callose accumulates at PDs along the longitudinal walls than at PDs along the transverse walls, as it happens in epidermal cells underlying the midvein (Gao et al., 2020). In the future, it will be interesting to distinguish between these possibilities; however, the mechanism by which changes in PD permeability are brought about during leaf development is inconsequential to the conclusions we derive from such changes.

### Auxin, Regulated PD Aperture, and Vein Patterning

Our results suggest that auxin controls PD permeability and that regulated PD aperture controls auxin-induced vein formation. Auxin application delays the reduction in PD permeability between veins and surrounding nonvascular tissues that occurs during normal leaf development. And in the most severe cases, impaired ability to regulate PD aperture almost entirely prevents auxin-induced vein formation.

Our results suggest that also auxin signaling and regulated PD aperture control each other during vein patterning. Auxin signaling inhibition prematurely reduces PD permeability between veins and surrounding nonvascular tissues (Fig. 7B), suggesting that auxin signaling normally delays such reduction. In turn, defects in the ability to regulate PD aperture lead to defects in expression of auxin response reporters (Fig. 7C). Near-constitutively narrow PD aperture leads to lower levels and broader domains of expression of auxin response reporters, suggesting that an auxin signal is produced at low levels in all cells and reaches veins through PDs. Also near-constitutively wide PD aperture leads to lower levels and broader domains of expression of auxin response reporters, suggesting that high levels of an auxin signal are maintained at sites of vein formation by reducing its PD-mediated leakage toward surrounding nonvascular tissues.

Our findings are consistent with the inability of plants with impaired ability to regulate PD aperture to restrict expression domains and maintain high expression levels of auxin response reporters in hypocotyl and root (Han et al., 2014; Liu et al., 2017; Mellor et al., 2020). Our interpretation is consistent with high levels of auxin signaling at early stages of vein formation and low levels of auxin signaling at late stages of vein formation (Mattsson et al., 2003; Krishna et al., 2021). And mutual control of auxin signaling and PD aperture regulation is consistent with the finding that simultaneous inhibition of auxin signaling and of the ability to regulate PD aperture leads to vein patterning defects that are more severe than the addition of the defects induced by auxin signaling inhibition and those induced by impaired ability to regulate PD aperture. In the most severe cases, simultaneous inhibition of auxin signaling and of the ability to regulate PD aperture leads to vascular systems comprised of very few, scattered vascular clusters.

It is unclear how auxin and its signaling could delay the reduction in PD permeability between veins and surrounding nonvascular tissues that occurs during normal leaf development. One possibility is that such delay is brought about by the ability of auxin to rapidly induce the expression of PD beta glucanases (Parizot et al., 2010; Benitez-Alfonso et al., 2013), which degrade callose at PDs and thus prevent callose-mediated restriction of PD aperture (Iglesias and Meins Jr, 2000; Levy et al., 2007; Rinne et al., 2011; Benitez-Alfonso et al., 2013). Another possibility is that the delay derives from the induction by auxin of pectin methylesterase activity (Bryan and Newcomb, 1954), which localizes around PDs (Morvan et al., 1998) and can increase their permeability (Chen et al., 2000; Chen and Citovsky, 2003; Dorokhov et al., 2012; Lionetti et al., 2014). A further possibility rests on the ability of auxin to reduce levels of reactive oxygen species in plastids (George et al., 2010), which leads to increased PD permeability (Benitez-Alfonso et al., 2009; Stonebloom et al., 2012). In the future, it will be interesting to test these possibilities; nevertheless, the mechanism by which auxin and its signaling delay the reduction in PD permeability between veins and surrounding nonvascular tissues has no bearing on our interpretation of such delay.

Our results also suggest that polar auxin transport and regulated PD aperture control each other during vein patterning. Auxin transport inhibition delays the reduction in PD permeability that occurs between veins and surrounding nonvascular tissues during normal leaf development (Fig. 7D), suggesting that polar auxin transport normally promotes such reduction. In turn, defects in the ability to regulate PD aperture lead to defects in expression and polar localization of the PIN1 auxin exporter (Fig. 7C), whose function is nonredundantly required for vein patterning (Sawchuk et al., 2013). Impaired ability to regulate PD aperture leads first to heterogeneous PIN1 expression along continuous expression domains and then to disconnection of those expression domains from pre-existing veins and breakdown of expression domains into domain fragments, suggesting that connection and continuity of PIN1 expression domains depend on regulated PD aperture. These defects in PIN1 expression resemble those of mutants in pathways that counteract *GN* function (Koizumi et al., 2005; Scarpella et al., 2006; Sieburth et al., 2006; Naramoto et al., 2009). And as in those mutants, in mutants impaired in PD aperture regulation PIN1 polarity is directed away from the edge of vein fragments and vascular clusters. That defects in the ability to regulate PD aperture lead to defects in PIN1 expression and localization is consistent with reduced polar auxin transport and defective PIN2 expression and localization in mutants and transgenics that are impaired in PD aperture regulation (Wu et al., 2016; Gao et al., 2020).

How polar auxin transport and regulated PD aperture could control each other during vein patterning remains to be explored, but PDs are associated with receptor-like proteins (Faulkner et al., 2013; Stahl et al., 2013; Vaddepalli et al., 2014) and PIN proteins with leucin-rich-repeat receptor kinases (Wang et al., 2013; Hajný et al., 2020), suggesting possibilities for the two pathways to interact. The molecular details of such interaction will have to be addressed in future research; however, our conclusion that polar auxin transport promotes the reduction in PD permeability that occurs between veins and surrounding nonvascular tissues is consistent with lower expression levels of positive regulators of callose production in auxin-transport-inhibited lateral roots (Sager et al., 2020). Moreover, mutual control of polar auxin transport and PD aperture regulation is consistent with the finding that simultaneous inhibition of auxin transport and the ability to regulate PD aperture leads to vein patterning defects that are more severe than the addition of the defects induced by auxin transport inhibition and those induced by impaired ability to regulate PD aperture. In the most severe cases, simultaneous inhibition of auxin transport and of the ability to regulate PD aperture leads to a vascularization zone that spans almost the entire width of the leaf. However, in those leaves veins still form oriented along the longitudinal axis of the leaf, suggesting the presence of residual vein patterning activity. That such residual vein patterning activity is provided by auxin signaling is suggested by the finding that the vascular system of leaves in which auxin signaling, polar auxin transport, and the ability to regulate PD aperture are simultaneously inhibited is no more than a shapeless cluster of vascular cells. This finding also suggests that those three pathways collectively provide all the vein patterning activity that exists in the leaf.

### Control of PD Aperture Regulation by GN

The vein pattern of leaves both lacking *GN* function and impaired in the ability to regulate PD aperture is no different from that of *gn* mutants. This suggests that *GN* controls PD aperture regulation, just as it controls auxin signaling and polar auxin transport (Verna et al., 2019). How *GN* controls PD aperture regulation is unclear, but the most parsimonious account is that *GN* controls the localization of proteins that regulate PD aperture. This hypothesis remains to be tested but is consistent with abnormal callose accumulation upon genetic or chemical inhibition of *GN* (Nielsen et al., 2012).

Irrespective of how *GN* precisely controls PD aperture regulation, simultaneous inhibition of auxin signaling, polar auxin transport, and the ability to regulate PD aperture phenocopies even the most severe vein patterning defects of *gn* mutants (Fig. 7E). Because vein patterning is prevented in both the strongest *gn* mutants and in the most severe instances of inhibition of auxin signaling, polar auxin transport, and the ability to regulate PD aperture, we conclude that vein patterning is the result of the coordinated action of three *GN*-dependent pathways: auxin signaling, polar auxin transport, and regulated PD aperture (Fig. 7F).

### A Diffusion–Transport-Based Vein-Patterning Mechanism

The Canalization Hypothesis was proposed more than 50 years ago to account for the inductive effects of auxin on vein formation (Sachs, 1968b; Sachs, 1981). In its most recent formulation (Sachs, 2000), the hypothesis proposes positive feedback between cellular auxin efflux mediated by exporters localized to a plasma membrane segment and localization of those auxin exporters to that membrane segment. The Canalization Hypothesis is supported by overwhelming experimental evidence and computational simulations; nevertheless, both experiments and simulations have brought to light inconsistencies between hypothesis and evidence (recently reviewed in (Ravichandran et al., 2020; Cieslak et al., 2021)). For example, the hypothesis assumes that at early stages of auxin-induced vein formation auxin diffuses from auxin sources (e.g., the applied auxin) toward auxin sinks (i.e. the pre-existing veins) (Sachs, 1981), but auxin diffusion out of the cell is unfavored over diffusion into the cell by almost two orders of magnitude (Runions et al., 2014). Furthermore, the hypothesis assumes that the veins whose formation is induced by auxin readily connect to pre-existing veins (i.e. auxin sinks) — an assumption that is supported by experimental evidence (Sachs, 1968a) but that simulations have been unable to reproduce without the addition of ad hoc solutions (Bayer et al., 2009; Smith and Bayer, 2009) or the existence of multiple auxin exporters with distinct patterns of auxin-responsive expression and polarization (O’Connor et al., 2014). Finally, the hypothesis relies on the function of auxin exporters (Sachs, 1984) — a requirement that is unsupported by experimental evidence because genetic or pharmacological inhibition of all the *PIN* genes with vascular function fails to prevent patterning of vascular cells into veins (Mattsson et al., 1999; Sieburth, 1999; Verna et al., 2019).

Our results suggest that those discrepancies between experiments and simulations, on the one hand, and the Canalization Hypothesis, on the other, could be resolved by supplementing the positive feedback between auxin and its polar transport postulated by the hypothesis with movement of an auxin signal through PDs according to its concentration gradient (Fig. 7A). At early stages of vein formation, movement through PDs would dominate; at later stages, polar transport would take over. Computational simulations suggests that our conclusion is justified (Cieslak et al., 2021).

A vein patterning mechanism that combines the positive feedback between auxin and its polar transport postulated by the Canalization Hypothesis with diffusion of an auxin signal through PDs requires at least two conditions to be met. First, auxin must promote the movement of an auxin signal through PDs such that gradual reduction in PD permeability between veins and surrounding nonvascular tissues as well as maintenance of symplastic connection between vein cells are accounted for by feedback between movement of the auxin signal through PDs and PD permeability. Our results support such requirement: auxin application delays the reduction in PD permeability that occurs during normal leaf development, thereby promoting movement of an auxin signal through PDs. Second, auxin signaling, polar auxin transport, and movement of an auxin signal through PDs must be coupled. Were they not — for example, if PD permeability between developing veins and surrounding nonvascular tissues remained high — the high levels of auxin signaling in early-stage PIN1 expression domains (Marcos and Berleth, 2014; Bhatia et al., 2019), which inefficiently transport auxin because of PIN1 isotropic localization (Benkova et al., 2003; Reinhardt et al., 2003; Heisler et al., 2005; Scarpella et al., 2006; Wenzel et al., 2007; Bayer et al., 2009; Sawchuk et al., 2013; Marcos and Berleth, 2014; Verna et al., 2015; Verna et al., 2019), would be dissipated by lateral diffusion of the auxin signal through PDs. And if, conversely, PD permeability in tissues where veins are forming was already low, the auxin signal would not be able to diffuse toward pre-existing veins, which transport auxin efficiently because of PIN1 polar localization and have low levels of auxin signaling (Benkova et al., 2003; Mattsson et al., 2003; Reinhardt et al., 2003; Heisler et al., 2005; Scarpella et al., 2006; Wenzel et al., 2007; Bayer et al., 2009; Sawchuk et al., 2013; Marcos and Berleth, 2014; Verna et al., 2015; Verna et al., 2019). That auxin signaling and polar auxin transport control each other during vein patterning is known (Mattsson et al., 2003; Wenzel et al., 2007; Donner et al., 2009; Krogan et al., 2012; Verna et al., 2015; Verna et al., 2019) (Fig. 7B,D). Our results support the additional requirement that polar auxin transport and movement of an auxin signal through PDs control each other, and that movement of an auxin signal through PDs and auxin signaling control each other (Fig. 7B–D).

In this study, we derived patterns of PD permeability change during leaf development from movement of a soluble YFP through leaf tissues. We note that auxin, being smaller than YFP, could for example move from the veins to the surrounding nonvascular tissues when YFP no longer can. Nevertheless, the reduced permeability of the PDs along the longitudinal walls of vein cells suggests that less auxin moves laterally than longitudinally. Moreover, unlike YFP, auxin is the substrate of PIN exporters (Petrasek et al., 2006; Zourelidou et al., 2014). By the time YFP can no longer move out of the veins, PIN1 has become polarly localized to the basal plasma membrane of vein cells (Benkova et al., 2003; Reinhardt et al., 2003; Heisler et al., 2005; Scarpella et al., 2006; Wenzel et al., 2007; Bayer et al., 2009; Sawchuk et al., 2013; Marcos and Berleth, 2014; Verna et al., 2015; Verna et al., 2019). Such polarization drives removal of auxin — but not YFP — from vein cells (Wisniewska et al., 2006), thereby dissipating the gradient in auxin signaling between veins and surrounding nonvascular tissues (Mattsson et al., 2003; Scarpella et al., 2006; Wenzel et al., 2007; Donner et al., 2009; Krogan et al., 2012; Marcos and Berleth, 2014; Verna et al., 2015; Verna et al., 2019), and as such the driving force for auxin movement from the veins to the surrounding nonvascular tissues. These considerations notwithstanding, the most stringent piece of evidence in support of our conclusions would be provided by the direct visualization of auxin movement. Despite considerable advances in the visualization of auxin signals (Brunoud et al., 2012; Liao et al., 2015; Herud-Sikimić et al., 2021), direct visualization of auxin movement remains to this day impossible. Should this change, it would also be possible to test whether it is auxin itself or an auxin-dependent signal that moves through PDs; nevertheless, the logic of the mechanism we report is unaffected by such limitation.

### Beyond Vein Patterning

Our observations pertain to vein patterning, but they may be relevant for other processes too — for example, stoma patterning. Indeed, like vein patterning, stoma patterning depends on auxin signaling (Balcerowicz et al., 2014; Le et al., 2014; Zhang et al., 2014), polar auxin transport (Le et al., 2014), regulated PD aperture (Guseman et al., 2010; Kong et al., 2012; Bundy et al., 2016), and *GN* (Mayer et al., 1993; Le et al., 2014). And as in vein patterning, stoma patterning defects in plants lacking *GN* function are quantitatively stronger than and qualitatively different from those in plants impaired in auxin signaling, polar auxin transport, or the ability to regulate PD aperture (Guseman et al., 2010; Kong et al., 2012; Balcerowicz et al., 2014; Le et al., 2014; Zhang et al., 2014; Bundy et al., 2016). However, whether the pathway network that patterns veins also patterns stomata remains to be tested.

Despite plants and animals acquired multicellularity independently of each other, a mechanism similar to that which patterns plant veins and depends on the movement of an auxin signal through PDs also patterns animal tissues. At early stages of tissue development in animal embryos, cells are connected by open gap junctions such that the tissue is a syncytium (reviewed in (Levin, 2007; Mathews and Levin, 2017)). And at later stages of tissue development, tissue compartments become isolated by selective closure of gap junctions to prevent unrestricted movement of signaling molecules. However, whereas in plants regulated PD aperture interacts with the polar transport of auxin and its signal transduction, in animals gap junction gating interacts with the polar secretion of signaling molecules or the binding of polarly localized ligands and receptors protruding from the plasma membranes (reviewed in (Levin, 2021)). It will be interesting to understand whether a mechanism that depends on the movement of signaling molecules through cell wall pores also patterns mushroom tissues; preliminary evidence (van Peer et al., 2010) suggests that it does.

## MATERIALS & METHODS

### Plants

Origin and nature of lines, genotyping strategies, and oligonucleotide sequences are in Table S1, Table S2, and Table S3, respectively. Seeds were sterilized and sowed as in (Sawchuk et al., 2008). Stratified seeds were germinated and seedlings were grown at 23 °C under continuous light (∼100 μmol m^−2^ s^−1^). Plants were grown at 24 °C under fluorescent light (∼100 μmol m^−2^ s^−1^) in a 16-h-light/8-h-dark cycle. Plants were transformed and representative lines were selected as in (Sawchuk et al., 2008).

### Chemicals

NPA and PBA were dissolved in dimethyl sulfoxide and stored at −20 °C indefinitely (NPA) or up to a week (PBA). Dissolved chemicals were added (25 μM final NPA concentration; 10 or 50 μM final PBA concentration) to growth medium just before sowing. IAA was dissolved in melted (55 C) lanolin (1% final IAA concentration) and stored at 4 C up to a week.

### Imaging

Developing leaves were mounted and imaged by confocal laser scanning microscopy as in (Sawchuk et al., 2013), except that emission was collected from ∼2.5–10-μm-thick optical slices. Mature leaves were fixed in 6 : 1 ethanol : acetic acid, rehydrated in 70% ethanol and water, cleared briefly (few seconds to few minutes) — when necessary — in 0.4 M sodium hydroxide, washed in water, and either (1) mounted in 1 : 2 : 8 water : glycerol : chloral hydrate and imaged by differential-interference-contrast or dark-field-illumination microscopy as in (Odat et al., 2014) or (2) stained for 6–16 h in 0.2% basic fuchsin in ClearSee (Kurihara et al., 2015), washed in ClearSee for 30 min, incubated in daily changed ClearSee for three days, and mounted in ClearSee for imaging by confocal laser scanning microscopy. Light paths for confocal laser scanning microscopy are in Table S4. In the Fiji distribution (Schindelin et al., 2012) of ImageJ (Schneider et al., 2012; Schindelin et al., 2015; Rueden et al., 2017), grayscaled RGB color images were turned into 8-bit images; when necessary, 8-bit images were combined into stacks, and stacks were projected at maximum intensity; look-up-tables were applied to images; and image brightness and contrast were adjusted by linear stretching of the histogram.

## ACKNOWLEDGEMENTS

We thank the Arabidopsis Biological Resource Center for *gsl8-6*/SAIL_679_H10, *gsl8-*1/SALK_111094, and *gsl8-2*/GK_851C04; Eva Benková and Jiří Friml for PIN1::PIN1:GFP; Jian Xu and Ben Scheres for PIN1::PIN1:YFP; Keiko Torii for *gsl8-chor*; Marcus Heisler and Elliot Meyerowitz for DR5rev::nYFP^HS^; Nico De Storme and Danny Geelen for *gsl8-et2*; and Yka Helariutta for *cals3-2d* and *casl3-3d*. We thank Przemek Prusinekiwcz, Mik Cieslak, and Adam Runions for insightful discussions. This work was supported by Discovery Grants of the Natural Sciences and Engineering Research Council of Canada to ES. NML was supported, in part, by a Summer Undergraduate Research Fellowship from the American Society of Plant Biologists.

