## Supplemental Tables for "Control of Leaf Vein Patterning by Regulated Plasmodesma Aperture"

**Table S1. Origin and Nature of Lines**

| *Line* | *Origin/Nature* |
|---|---|
| *cals3-2d* | (Vatén et al., 2011); introgressed into Col-0 |
| *cals3-3d* | (Vatén et al., 2011) |
| *gsl8-et2* | (De Storme et al., 2013) |
| *gsl8-6* | SAIL_679_H10 (ABRC); (Chen et al., 2009; Sessions et al., 2002) |
| *gsl8-chor* | (Guseman et al., 2010) |
| *gsl8-1* | SALK_111094 (ABRC); (Alonso et al., 2003; Töller et al., 2008) |
| *gsl8-2* | GK_851C04 (ABRC); (Kleinboelting et al., 2012; Töller et al., 2008) |
| UAS::YFP | Transcriptional fusion of six copies of the UAS sequence (Giniger et al., 1985) upstream of the -46 Cauliflower Mosaic Virus 35S promoter (Odell et al., 1985) to a translationally enhanced Venus-encoding sequence (Gallie et al., 1988; Nagai et al., 2002) (primers: "TeVENUS Fwd XbaI" and "TeVENUS Rev SacI") |
| E2331 | (Amalraj et al., 2020; Gardner et al., 2009) |
| Q0990 | (Haseloff, 1999; Sawchuk et al., 2007) |
| Q0950 | (Haseloff, 1999; Sawchuk et al., 2007) |
| J3281 | (Haseloff, 1999; Sawchuk et al., 2007) |
| J1701 | (Haseloff, 1999; Sawchuk et al., 2007) |
| PIN1::PIN1:YFP | (Xu et al., 2006) |

| *Line* | *Origin/Nature* |
| --- | --- |
| PIN1::PIN1:GFP | (Benková et al., 2003) |
| DR5rev::nYFP$^{HS}$ | (Heisler et al., 2005; Sawchuk et al., 2013) |
| DR5rev::nYFP$^{ES}$ | Transcriptional fusion of nine copies of the DR5Rev sequence (Ulmasov et al., 1997) upstream of the -46 Cauliflower Mosaic Virus 35S promoter (Odell et al., 1985) to EYFP-Nuc (Clontech) |
| *gn-13* | (Alonso et al., 2003; Verna et al., 2019) |

*Supplemental References*


**Alonso, J. M., Stepanova, A. N., Leisse, T. J., Kim, C. J., Chen, H., Shinn, P., Stevenson, D. K., Zimmerman, J., Barajas, P., Cheuk, R., et al.** (2003). Genome-wide insertional mutagenesis of Arabidopsis thaliana. *Science (80-. ).* **301**, 653–657.

**Amalraj, B., Govindaraju, P., Krishna, A., Lavania, D., Linh, N. M., Ravichandran, S. J. and Scarpella, E.** (2020). GAL4/GFP enhancer-trap lines for identification and manipulation of cells and tissues in developing Arabidopsis leaves. *Dev. Dyn.* **249**, 1127–1146.

**Benková, E., Michniewicz, M., Sauer, M., Teichmann, T., Seifertová, D., Jürgens, G. and Friml, J.** (2003). Local, efflux-dependent auxin gradients as a common module for plant organ formation. *Cell* **115**, 591–602.

**Chen, X.-Y., Liu, L., Lee, E., Han, X., Rim, Y., Chu, H., Kim, S.-W., Sack, F. and Kim, J.-Y.** (2009). The Arabidopsis callose synthase gene GSL8 is required for cytokinesis and cell patterning. *Plant Physiol.* **150**, 105–113.

**De Storme, N., De Schrijver, J., Van Criekinge, W., Wewer, V., Dörmann, P. and Geelen, D.** (2013). GLUCAN SYNTHASE-LIKE8 and STEROL METHYLTRANSFERASE2 are required for ploidy consistency of the sexual reproduction system in Arabidopsis. *Plant*

*Cell* **25**, 387–403.

**Gallie, D. R., Sleat, D. E., Watts, J. W., Turner, P. C. and Wilson, T. M.** (1988). Mutational analysis of the tobacco mosaic virus 5′-leader for altered ability to enhance translation. *Nucleic Acids Res.* **16**, 883–893.

**Gardner, M. J., Baker, A. J., Assie, J.-M., Poethig, R. S., Haseloff, J. P. and Webb, A. A. R.** (2009). GAL4 GFP enhancer trap lines for analysis of stomatal guard cell development and gene expression. *J. Exp. Bot.* **60**, 213–226.

**Giniger, E., Varnum, S. M. and Ptashne, M.** (1985). Specific DNA binding of GAL4, a positive regulatory protein of yeast. *Cell* **40**, 767–774.

**Guseman, J. M., Lee, J. S., Bogenschutz, N. L., Peterson, K. M., Virata, R. E., Xie, B., Kanaoka, M. M., Hong, Z. and Torii, K. U.** (2010). Dysregulation of cell-to-cell connectivity and stomatal patterning by loss-of-function mutation in Arabidopsis chorus (glucan synthase-like 8). *Development* **137**, 1731–1741.

**Haseloff, J.** (1999). Chapter 9: GFP Variants for Multispectral Imaging of Living Cells. In *Green Fluorescent Proteins* (ed. Sullivan, K. F.) and Kay, S. A. B. T.-M. in C. B.), pp. 139–151. Academic Press.

**Heisler, M. G., Ohno, C., Das, P., Sieber, P., Reddy, G. V., Long, J. A. and Meyerowitz, E. M.** (2005). Patterns of auxin transport and gene expression during primordium development revealed by live imaging of the Arabidopsis inflorescence meristem. *Curr. Biol.* **15**, 1899–1911.

**Kleinboelting, N., Huep, G., Kloetgen, A., Viehoever, P. and Weisshaar, B.** (2012). GABI-Kat SimpleSearch: new features of the Arabidopsis thaliana T-DNA mutant database. *Nucleic Acids Res.* **40**, D1211-15.

**Nagai, T., Ibata, K., Park, E. S., Kubota, M., Mikoshiba, K. and Miyawaki, A.** (2002). A variant of yellow fluorescent protein with fast and efficient maturation for cell-biological applications. *Nat. Biotechnol.* **20**, 87–90.

**Odell, J. T., Nagy, F. and Chua, N. H.** (1985). Identification of DNA sequences required for activity of the cauliflower mosaic virus 35S promoter. *Nature* **313**, 810–812.

**Sawchuk, M. G., Head, P., Donner, T. J. and Scarpella, E.** (2007). Time-lapse imaging of Arabidopsis leaf development shows dynamic patterns of procambium formation. *New Phytol.* **176**, 560–571.

**Sawchuk, M. G., Edgar, A. and Scarpella, E.** (2013). Patterning of leaf vein networks by convergent auxin transport pathways. *PLoS Genet.* **9**, e1003294.

**Sessions, A., Burke, E., Presting, G., Aux, G., McElver, J., Patton, D., Dietrich, B., Ho, P., Bacwaden, J., Ko, C., et al.** (2002). A high-throughput Arabidopsis reverse genetics system. *Plant Cell* **14**, 2985–2994.

**Töller, A., Brownfield, L., Neu, C., Twell, D. and Schulze-Lefert, P.** (2008). Dual function of Arabidopsis glucan synthase-like genes GSL8 and GSL10 in male gametophyte development and plant growth. *Plant J.* **54**, 911–923.

**Ulmasov, T., Murfett, J., Hagen, G. and Guilfoyle, T. J.** (1997). Aux/IAA proteins repress expression of reporter genes containing natural and highly active synthetic auxin response elements. *Plant Cell* **9**, 1963–1971.

**Vatén, A., Dettmer, J., Wu, S., Stierhof, Y.-D., Miyashima, S., Yadav, S. R., Roberts, C. J., Campilho, A., Bulone, V., Lichtenberger, R., et al.** (2011). Callose biosynthesis regulates symplastic trafficking during root development. *Dev. Cell* **21**, 1144–1155.

**Verna, C., Ravichandran, S. J., Sawchuk, M. G., Linh, N. M. and Scarpella, E.** (2019). Coordination of tissue cell polarity by auxin transport and signaling. *Elife* **8**, 1–30.

**Xu, J., Hofhuis, H., Heidstra, R., Sauer, M., Friml, J. and Scheres, B.** (2006). A molecular framework for plant regeneration. *Science* **311**, 385–388.

**Table S2. Genotyping Strategies**

| *Line* | *Genotyping Strategy* |
|---|---|
| *cals3-2d* | "CALS3 FWD 1" and "CALS3 m REV2" |
| *cals3-3d* | "cals3-3d F" and "cals3-3d R"; *Taq*I |
| *gsl8-et2* | "ET2dCAPS F" and "ET2dCAPS R2"; *Hin*dIII |
| *gsl8-6* | *GSL8*: "SAIL_679_H10LP" and "SAIL_679_H10RP"; *gsl8-6*: "SAIL_679_H10RP" and "LBb1.3" |
| *gsl8-chor* | "chorus dCAPS F" and "chorus dCAPS R"; *Nla*IV |
| *gsl8-1* | *GSL8*: "GSL8 FWD" and "GSL8 REV"; *gsl8-1*: "GSL8 FWD" and "LBb1.3" |
| *gsl8-2* | *GSL8*: "GK_851C04LP" and "GK_851C04RP"; *gsl8-2*: "GK_851C04RP" and "08474" |
| *gn-13* | *GN*: "SALK_045424 gn LP" and "SALK_045424 gn RP"; *gn*: "SALK_045424 gn RP" and "LBb1.3" |

**Table S3. Oligonucleotide Sequences**

| *Name* | *Sequence (5′ to 3′)* |
|---|---|
| CALS3 FWD 1 | ATCCCTTGTCAACTCAGG |
| CALS3 m REV2 | GAGAGATCTGAAGAGCTT |
| cals3-3d F | CCATCTCTTGTGCAACTTTACAATG |
| cals3-3d R | TATCAGGATCGAGAGGTAGGATATTATCG |
| ET2dCAPS F | AAATTGTATTGGATCGTGACCTGTAATCTTTCATGC |
| ET2dCAPS R2 | CTCCAATATCCTTCCTCTTAATGTTTGGATATAAGC |
| SAIL_679_H10LP | GCCCAGGTATACTAAGCTGGG |
| SAIL_679_H10RP | CTTTTTCTTCTAACGTGGGGG |
| LBb1.3 | ATTTTGCCGATTTCGGAAC |
| chorus dCAPS F | TCATGTGGATGCTTAGTGAACTGCTTCTTACTAACT |
| chorus dCAPS R | AGCCAACTGACCCAGTCTTCAAAATCCTCGAGGGTC |
| GSL8 FWD | TCACATGCATATAGCTGTGGG |
| GSL8 REV | TAGTTCCGCAGACAAAGTTGC |
| GK_851C04LP | TTCAGAAGTTGCATCTGCATG |
| GK_851C04RP | ACACTCTGGAAGAAAGCGGAC |
| 08474 | ATAATAACGCTGCGGACATCTACATTTT |

| *Name* | *Sequence (5′ to 3′)* |
|---|---|
| TeVENUS Fwd XbaI | GCGCGCTCTAGAGTATTTTTACAACAATTACCAACAACAAC |
| TeVENUS Rev SacI | AAAGAGCTCTTACTCGTCCATGCCGAGAGTG |
| SALK_045424 gn LP | TGATCCAAATCACTGGGTTTC |
| SALK_045424 gn RP | AGCTGAAGATAGGGAATTCGC |

**Table S4. Confocal Light Paths**

| *Fluorophore* | *Laser* | *Wavelength (nm)* | *Main Dichroic Beam Splitter* | *First Secondary Dichroic Beam Splitter* | *Second Secondary Dichroic Beam Splitter* | *Emission Filter (Detector)* |
|---|---|---|---|---|---|---|
| Lignin | HeNe | 543 | HFT 405/488/543 | Mirror | NFT 515 | BP 600–650 (PMT3) |
| YFP; Autofluorescence | Ar | 514 | HFT 405/514/594 | NFT 595 | NFT 515 (PMT3); Plate (META) | BP 520-555 IR (PMT3); 593–754 (META) |
| GFP; YFP | Ar | 458; 514 | HFT 458/514 | NFT 595 | NFT 545 (PMT2); NFT 545 (PMT3) | BP 475–525 (PMT2); BP 520-555 IR (PMT3) |
| GFP; YFP; Autofluorescence | Ar | 458; 514 | HFT 458/514 | NFT 595 | NFT 545 (PMT2); NFT 545 (PMT3); Plate (META) | BP 475–525 (PMT2); BP 520-555 IR (PMT3); 657–754 (META) |
| GFP; Autofluorescence | Ar | 488 | HFT 405/488/594 | NFT 545 | NFT 490 (PMT3); Plate (META) | BP 505–530 (PMT3); 550–754 (META) |
| YFP | Ar | 514 | HFT 405/514/594 | NFT 595 | NFT 515 | BP 520-555 IR (PMT3) |
